# Organophosphate intoxication in *C. elegans* reveals a new route to mitigate poisoning through the modulation of determinants responsible for nicotinic acetylcholine receptor function

**DOI:** 10.1101/2021.05.01.442241

**Authors:** Patricia G. Izquierdo, Claude L. Charvet, Cedric Neveu, A. Christopher Green, John E.H. Tattersall, Lindy Holden-Dye, Vincent O’Connor

**Affiliations:** Biological Sciences, Institute for Life Sciences, University of Southampton, Southampton, SO17 1BJ, United Kingdom; French National Institute for Agricultural Research (INRA), Infectiologie Animale et Santé Publique, Nouzilly, France; Dstl, Defence Science and Technology Laboratory, Porton Down, Salisbury, Wiltshire, SP4 0JQ, United Kingdom

**Keywords:** Carbamate, organophosphate, nicotinic receptor, aldicarb, paraoxon, oxime, neuromuscular junction, cholinergic plasticity, plasticity-promoting treatment

## Abstract

Plasticity is a reactive mechanism that allows the adaptation of organisms to changing environmental cues. The exploitation of this physiological process has a clear benefit to promote the recovery from a wide range of neurological disorders. Here, we show that plasticity-promoting regimes provide candidate mechanisms to supplement the classically used antidotes for anti-cholinesterase poisoning. These neurotoxins inhibit acetylcholinesterase, causing the overstimulation of cholinergic transmission at synapses and neuromuscular junctions. The model organism *C. elegans* exhibits organophosphate-induced mitigating plasticity that impacts on the recovery of neuromuscular phenotypes, initially impaired by the drug. This is underpinned by overstimulation of nicotinic receptors at the neuromuscular junction. Intrinsic determinants of receptor’s location and sensitivity modulate the extent of plasticity in the context of persistent cholinergic stimulation. Our results indicate that pharmacological intervention of nicotinic receptors and/or scaffolding proteins that support receptor function might provide a novel treatment route for anti-cholinesterase poisoning.

## Introduction

Molecular plasticity is a conserved mechanism from invertebrates to mammals involved in the adaptation and survival of the living organisms. This principle requires the reorganization of a signalling cascade in order to modulate the strength of output responses to intrinsic or extrinsic stimuli. (Citri and Malenka 2008, Markram, Gerstner et al. 2011, Mateos-Aparicio and Rodriguez-Moreno 2019). The expression of such molecular mechanism is fundamental to neural plasticity and the reactive changes associated with distinct forms of synaptic homeostasis (Thompson 1986, Okano, Hirano et al. 2000, Byrne, LaBar et al. 2014). Pharmacological modulation of naturally occurring plasticity could be beneficial in treating damage in the nervous system. This idea could be extended to conditions in which neurotoxins poisoning impairs neurotransmitter signalling. Acetylcholinesterase inhibitors, organophosphates and carbamates, are two important class of such neurotoxins. Anti-cholinesterase poisoning raises the concentration of acetylcholine at the synapses beyond the physiological levels and triggers the persistent stimulation of the nicotinic and muscarinic receptors (Colovic, Krstic et al. 2013, Tattersall 2018). Since acetylcholine regulates transmission at central, peripheral and neuromuscular synapses (Koelle 1954, Massoulie, Pezzementi et al. 1993), there is a wide spectrum of symptoms associated with anti-cholinesterase poisoning, known as cholinergic syndrome (Jokanovic and Kosanovic 2010, Tattersall 2018). Asphyxia is the main cause of death and is produced by the uncontrolled stimulation of the nicotinic receptors at the neuromuscular junction of respiratory muscles (Jokanovic and Kosanovic 2010). The core of the treatment consists of the artificial ventilation of the victim and the injection of atropine, an antagonist of the muscarinic receptors (Eddleston and Chowdhury 2016). It is usually supplemented with an oxime treatment to facilitate reactivation of the acetylcholinesterase and an anticonvulsant drug to minimize seizures during the initial cholinergic crisis (Eddleston, Szinicz et al. 2002, Jokanovic and Prostran 2009, Eddleston and Chowdhury 2016, Jokanovic and Petrovic 2016). However, the success of this treatment depends on factors such as the time of reaction between organophosphate and acetylcholinesterase, the dose of atropine administrated or the type of cholinesterase inhibitor intoxicating (Eddleston, Buckley et al. 2004, Eddleston and Chowdhury 2016, Worek, Thiermann et al. 2016). In this scenario, developing alternative treatment strategies remains pressing (Jeyaratnam 1990, Konradsen 2007).

The cholinergic signalling, whether organizing central or peripheral transmission, exhibits highly plastic mechanisms that modulate the strength of the acetylcholine signalling. Structural changes in the cholinergic circuit have been demonstrated after a brain injury and this has an implication in the recovery from symptoms observed after rehabilitation (Wang, Conner et al. 2016). Cholinergic receptors are an additional checkpoint to regulate the level of acetylcholine signalling.

Fast cholinergic synaptic transmission is mediated by nicotinic receptors, acetylcholine-gated cation channels formed by five subunits in homomeric or heteromeric combination (Gotti, Fornasari et al. 1997). Since multiple genes encode for nicotinic receptor subunits, the combinatorial complexity might provide another opportunity to modulate the cholinergic signal (Gotti, Fornasari et al. 1997). Receptor subtypes exhibit different biophysical properties and some subtypes can be predominately expressed respect to others depending on external stimuli (Corriveau, Romano et al. 1995, Missias, Chu et al. 1996, Romano, Pugh et al. 1997). Pharmacological activation of nicotinic receptors by exogenous agonists like nicotine is an established route to modulation of receptor composition and function (Marks, Burch et al. 1983, Schwartz and Kellar 1983, Fenster, Whitworth et al. 1999, Nashmi, Xiao et al. 2007). Finally, the interaction of nicotinic receptors with auxiliary proteins modifies their trafficking, clustering, sensitivity or motility between synaptic and extrasynaptic domains with profound impacts on the overlying efficacy of cholinergic transmission. (Jeanclos, Lin et al. 2001, Sanes and Lichtman 2001, Lin, Jeanclos et al. 2002, Lansdell, Gee et al. 2005, Boulin, Rapti et al. 2012, Miwa, Anderson et al. 2019). Understanding the molecular mechanisms that regulate the cholinergic signal at all levels would be critical to develop plasticity-promoting treatments in the context of anti-cholinesterase intoxication.

Despite its simplicity, the free-living nematode *C. elegans* can develop behavioural plasticity by two different paradigms known as associative and non-associative (Hobert 2003, Ardiel and Rankin 2010). In particular, preconditioning of nematodes to nicotine and other environmental cues such as starvation or temperature modifies the consequent phenotype of these worms when they are post-exposed to the same signal (Hedgecock and Russell 1975, Mori 1999, Sawin, Ranganathan et al. 2000, Feng, Li et al. 2006). Alternatively, the chronic exposure to nicotine triggers the habituation of egg-laying, the cholinergic-dependent behaviour responsible for reproduction (Waggoner, Dickinson et al. 2000, Smith, Zhang et al. 2013). The genetic amenability of the nematode combined with a well-defined set of behaviours and the characterization of its nervous system make *C. elegans* an attractive model organism to research the genetics of behavioural plasticity including those associated with cholinergic transmission (Waggoner, Dickinson et al. 2000, Hobert 2003, Feng, Li et al. 2006, Smith, Zhang et al. 2013).

In previous experiments, we built on the established work of others and demonstrated the potential of the organism model *C. elegans* to investigate acetylcholinesterase intoxication and recovery (Izquierdo, O’Connor et al. 2020). Specifically, we highlighted the measurement of pumping rate on food as a suitable cholinergic-meadiated behaviour that is dose-time dependent inhibited by the presence of anti-cholinesterases but can be restored when nematodes are removed from the drug condition (Izquierdo, O’Connor et al. 2020). Here, we demonstrate that nematodes express a cholinergic plasticity in two different paradigms, the precondition to low doses of the drug and the chronic stimulation with high intoxicating concentrations. The preconditioning paradigm with the organophosphate paraoxon-ethyl intensified the behavioural effect of the drug when nematodes are post-exposed. However, the incubation to large concentrations of paraoxon-ethyl triggers a mitigating behavioural plasticity observed by the spontaneous recovery of the cholinergic-dependent pharyngeal pumping that underlies feeding. We identified mutants that impact the synaptic organization and/or sensitivity of the nicotinic receptors at the neuromuscular junction as important determinants controlling the scale of plasticity in a positive or negative manner. This finding provokes the notion that new approaches can be used to modulate the tone of the nicotinic acetylcholine receptor function during organophosphate intoxication. Importantly, this modulation might mitigate the effects of the drug poisoning opening new insights to develop alternative strategies to treat anti-cholinesterase intoxication.

## Results

### The pharyngeal microcircuit of *C. elegans* exhibits mitigating aldicarb-induced plasticity after preconditioning with sub-maximal dose of the drug

*C. elegans* is able to adapt to environmental conditions depending on their previous experience (Bernhard and van der Kooy 2000, Ardiel and Rankin 2010, Kunitomo, Sato et al. 2013). As a paradigm of behavioural plasticity in *C. elegans*, we investigated how the preconditioning to acetylcholinesterase inhibitors impacted on the subsequent response to drug treatment. This was done by quantifying pumping phenotype on food, a cholinergic-dependent behaviour that exhibits a dose-time dependent inhibition to the exposure of anti-cholinesterases (Izquierdo, O’Connor et al. 2020). We observed that nematodes exposed to 50 μM aldicarb for 24 hours exhibited a 50% drug-dependent inhibition of the pharyngeal function. However, recovery from inhibition was observed 2 hours after being removed from the drug (Fig. 1). This implies a reversible pharmacological inhibition of the enzyme without affecting the underpinning functions that control the tested behaviour.

**Figure 1.**
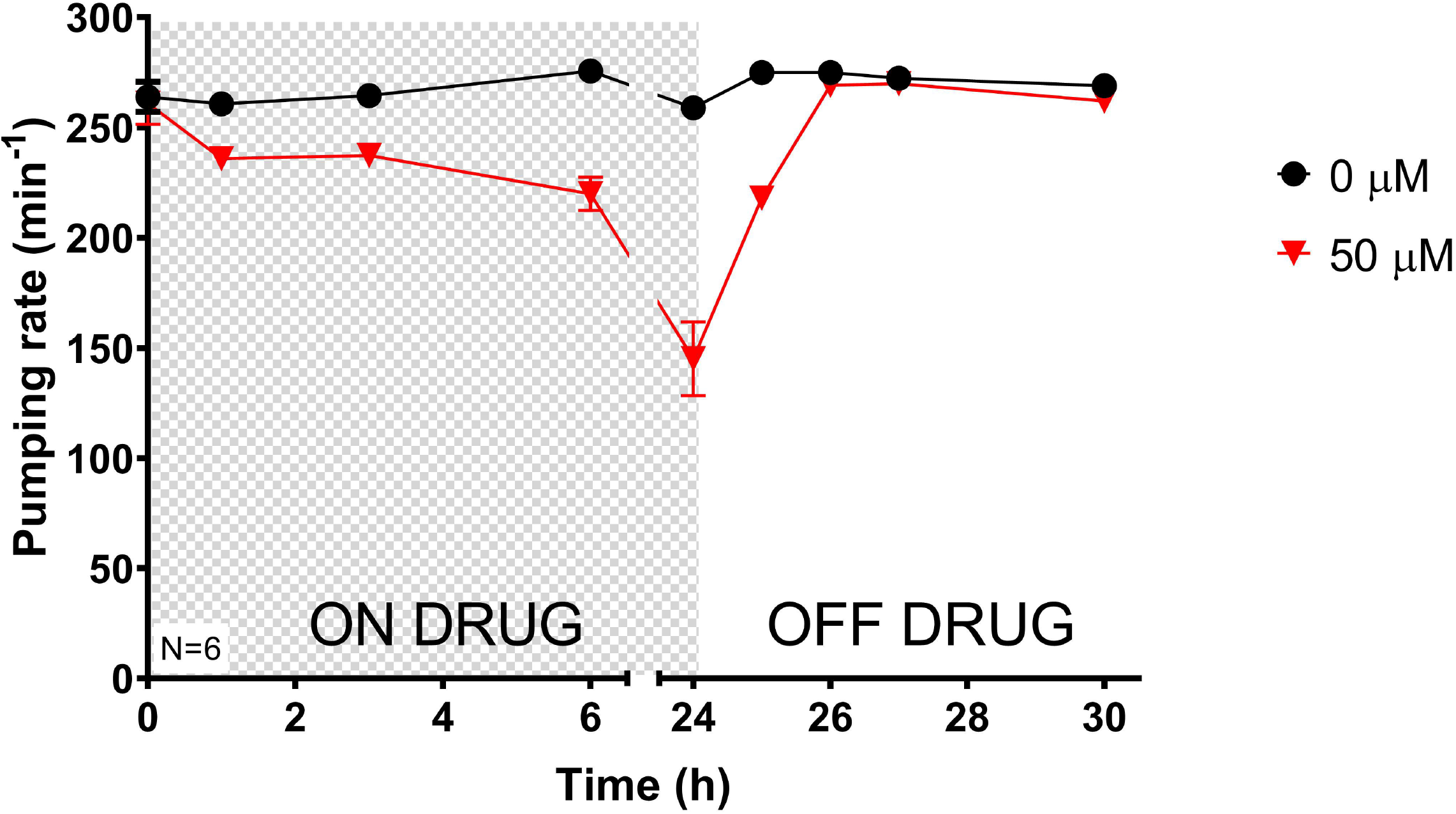
Worms intoxicated with sub-lethal dose of aldicarb exhibited recovery of the pharyngeal function after 2h of being removed from the drug-containing plate. Pharyngeal pumping rate per minute was quantified at different end-point times for synchronized L4+1 nematodes exposed to a sub-lethal dose of aldicarb. Nematodes were then transferred onto non-drug containing plates where the recovery of the pharyngeal function was observed within two hours of being removed from the drug. Shaded box indicates period of treatment. Data are shown as mean ± SEM of 6 worms in at least 3 independent experiments. Statistical significance between exposed and non-exposed nematodes was calculated by two-way ANOVA test followed by Bonferroni corrections. **p≤0.01; ***p≤0.001.

Based on the above, nematodes were incubated either in control plates or 50 μM aldicarb-containing plates for 24 hours and then transferred onto non-drugged plates for 2 hours to allow the recovery of the pharyngeal function. Finally, control and aldicarb-treated worms were intoxicated on plates containing fivefold increase of anti-cholinesterase concentration where pump rate on food was measured. (Fig. 2A). Wild type nematodes exposed to the maximal concentration of aldicarb exhibited a time-dependent inhibition of the feeding phenotype, being completely abolished after 24 hours of incubation (Izquierdo, O’Connor et al. 2020). Interestingly, the pharyngeal pumping rate of nematodes pre-exposed to the cholinesterase inhibitor aldicarb was less susceptible to inhibition than non-preconditioned worms when they were subsequently intoxicated with the maximal dose. This was clearly evidenced at 6 and 24 hours of being transferred to the 250 μM of aldicarb-containing plates (Fig. 2B).

**Figure 2.**
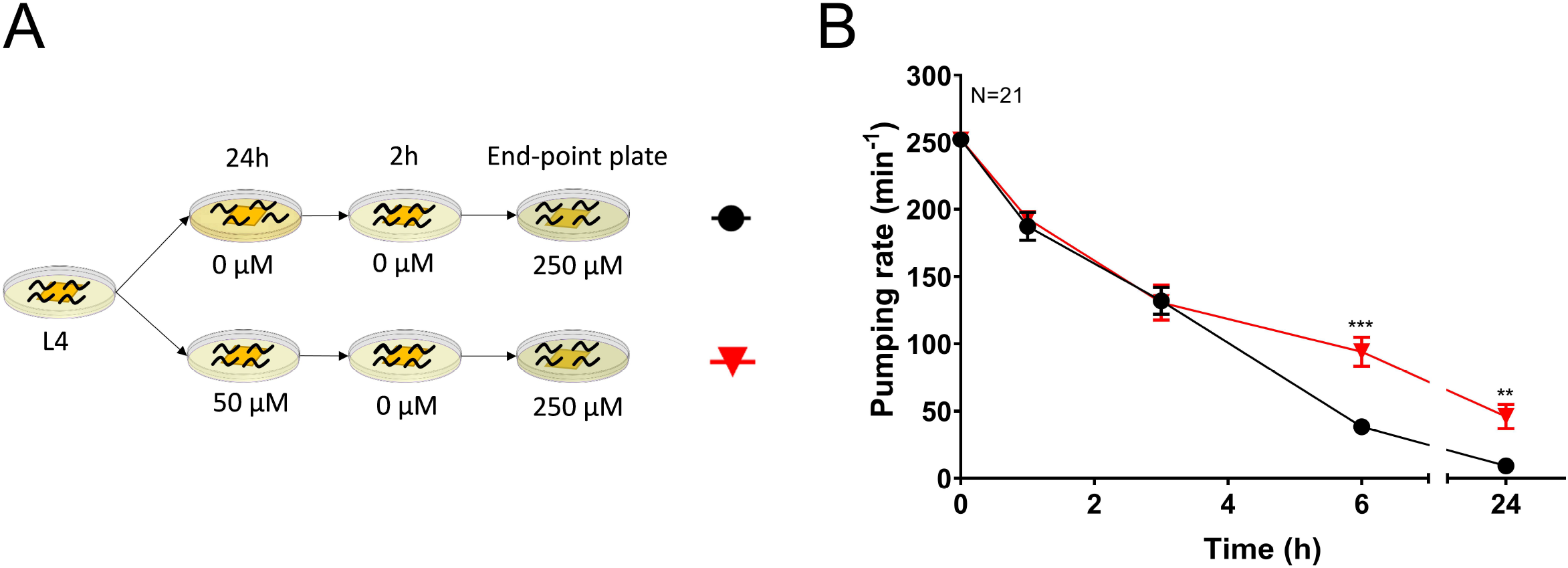
Nematodes preconditioned with aldicarb exhibit a late reduction in sensitivity of the pharyngeal circuits to the exposure of an increased dose of the drug. A) Synchronized L4 worms were intoxicated onto 50 μM aldicarb plates. Non-exposed nematodes were used as control. After 24 hours, they were transferred onto non-drug containing plates to allow the recovery of the pharyngeal function before exposing them to 250 μM aldicarb plates where the pharyngeal pumping rate was scored. B) Preconditioned worms exhibited higher pharyngeal pumping rate than non-preconditioned animals after 6 and 24 hours of being transferred onto the maximal dose plates. Data are shown as mean ± SEM of 21 worms in at least 11 independent experiments. Statistical significance between preconditioned and non-preconditioned nematodes was calculated by two-way ANOVA test followed by Bonferroni corrections. **p≤0.01; ***p≤0.001.

The result indicates that the preconditioning step with aldicarb induces a mitigating plasticity in the pharyngeal phenotype of pre-exposed worms when they are subsequently exposed to a maximal drug concentration.

### Preconditioning with sub-maximal dose of paraoxon-ethyl leads to an aggravating plasticity effect in the pharyngeal phenotype

Carbamates and organophosphates are two distinct groups of cholinesterase inhibitors that cause similar cholinergic toxicity (Colovic, Krstic et al. 2013). In order to investigate the behavioural plasticity of the feeding phenotype that emerges from organophosphate preconditioning, we performed an equivalent experiment to the above described with the organophosphate paraoxon-ethyl (Fig. 3A). We previously demonstrated that paraoxon-ethyl inhibits *C. elegans* acetylcholinesterases in a reversible manner (Izquierdo, O’Connor et al. 2020). This triggers an inhibition of the pharyngeal function that is recoverable when nematodes are removed from drugged plates (Izquierdo, O’Connor et al. 2020). In the present study, wild type nematodes were preconditioned for 24 hours on plates containing 20 μM of paraoxon-ethyl. Similar to aldicarb experiment, this concentration and time of exposure was selected to reduce the pumping phenotype by half of the maximal response after 24 hours of intoxication. The inhibition was recovered after 3 hours of being removed from the drugged plates. Finally, the preconditioned and non-preconditioned worms were transferred to 100 μM paraoxon-ethyl, a fivefold higher concentration than used in the preceding preconditioning step (Fig. 3A).

**Figure 3.**
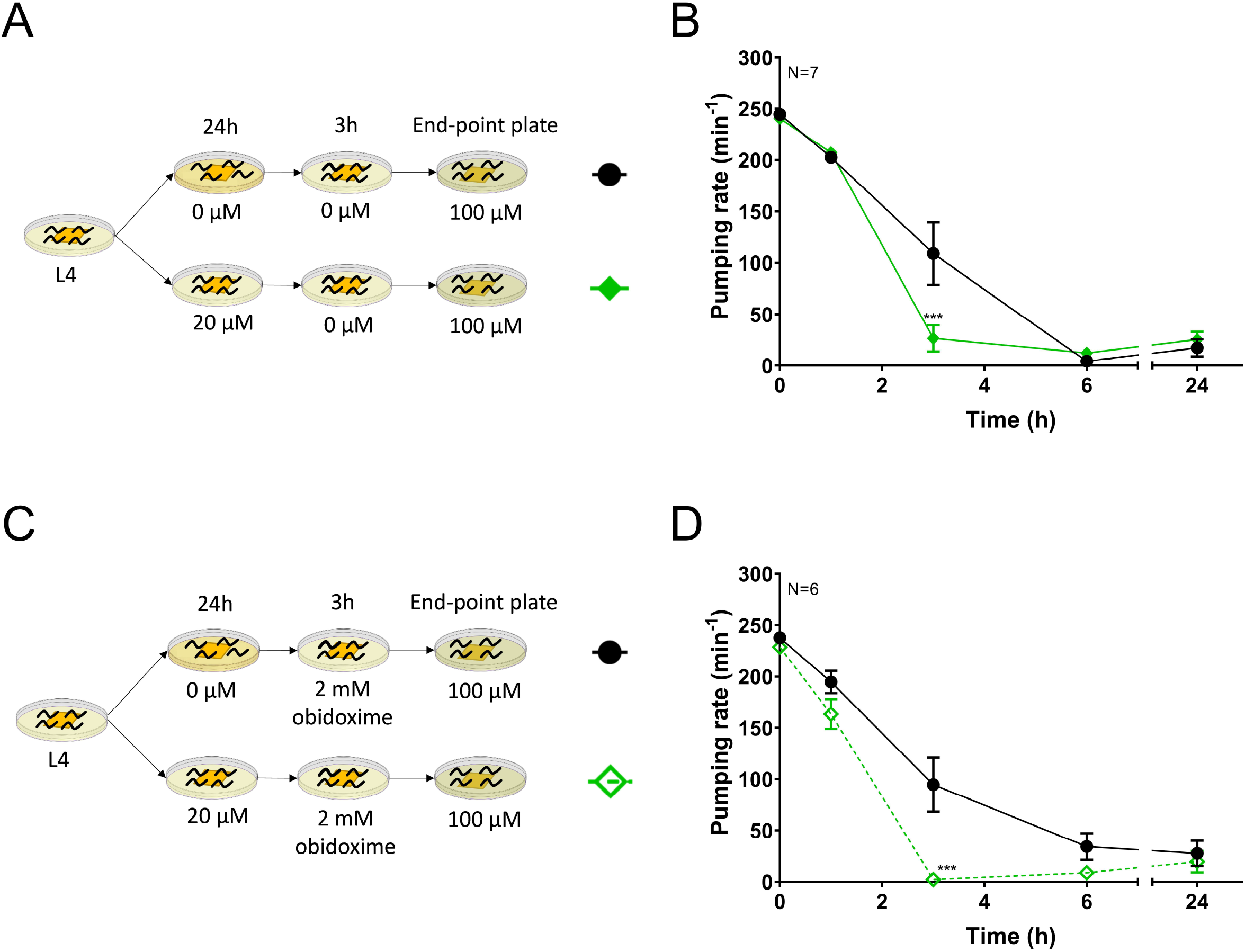
Nematodes preconditioned with paraoxon-ethyl are sensitized to subsequent OP inhibition of the pharyngeal pumping. A) Synchronized L4 worms were incubated onto either non-drug or 20 μM paraoxon-ethyl containing plates. After 24 hours, they were transferred onto non-drug containing plates to allow the recovery of the pharyngeal function. Following, they were picked onto 100 μM paraoxon-ethyl plates where the pharyngeal pumping was scored. B) OP- preconditioned nematodes exhibited a greater reduction of the pharyngeal pumping after 3h transfer to 100 μM paraoxon-ethyl plates compared to the non-preconditioned animals. Data are shown as mean ± SEM of 7 worms in at least 4 independent experiments. C) Nematodes were preconditioned as indicated in A, however, obidoxime was added to the recovery plate to promote the rescue of the acetylcholinesterase activity after paraoxon-ethyl inhibition. D) Preconditioned nematodes exhibited a similar response to maximal dose of paraoxon-ethyl when they were allowed to recover in the presence or in the absence of obidoxime. Data are shown as mean ± SEM of 6 worms in at least 3 independent experiment. Statistical significance between preconditioned and non-preconditioned nematodes was calculated by two-way ANOVA test followed by Bonferroni corrections. ***p≤0.001.

In contrast to the results observed for aldicarb, the pre-exposure to paraoxon-ethyl caused an aggravated inhibition of pumping compared to the control non-precondition treatment. This was evidenced by the residual pumping observed after 3 hours of transferring nematodes to the maximal concentration plates (Fig. 3B). This intensified inhibition could be explained by either an increased sensitivity to paraoxon-ethyl of preconditioned worms or an incomplete recovery of the acetylcholinesterase inhibition imposed by the initial exposure phase of the protocol. Recovery studies from other organophosphate pesticides intoxication demonstrate that acetylcholinesterase activity was incompletely recovered even when nematodes present a normal phenotype based on visual observations (Opperman and Chang 1991, Lewis, Gehman et al. 2013). In order to investigate this, we supplemented the recovery plate with 2 mM obidoxime (Fig. 3C). We previously demonstrated that obidoxime improves the acetylcholinesterase activity and the pharyngeal function of *C. elegans* during the recovery from paraoxon-ethyl inhibition (Izquierdo, O’Connor et al. 2020). Similar to previous observations, preconditioned nematodes remained more susceptible to the maximal dose of paraoxon-ethyl at 3 hours of incubation compared to those never exposed to the drug (Fig. 3D). This indicates that the aggravated behavioural plasticity observed in the preconditioned nematodes with paraoxon-ethyl is due to an increased sensitivity to the drug rather than a residual inhibition of the worm acetylcholinesterase.

The distinct pattern of plasticity observed between aldicarb and paraoxon-ethyl preconditioned worms suggests differences in the drug-dependent mechanism of adaptation to the acetylcholinesterase inhibition by carbamates and organophosphates despite sharing a core mode of action.

### Aldicarb preconditioning treatment avoid the synaptic plasticity of nematodes post-exposed to paraoxon-ethyl

In order to interrogate the nature of the distinct preconditioning outcomes with carbamates and organophosphates, nematodes were pre-exposed with aldicarb and then post-exposed to paraoxon-ethyl (Fig. 4A). Preconditioned and non-preconditioned nematodes to 50 μM aldicarb for 24 hours exhibited a similar inhibition pattern of the pumping rate when they were subsequently exposed to 100 μM paraoxon-ethyl (Fig. 4B).

**Figure 4.**
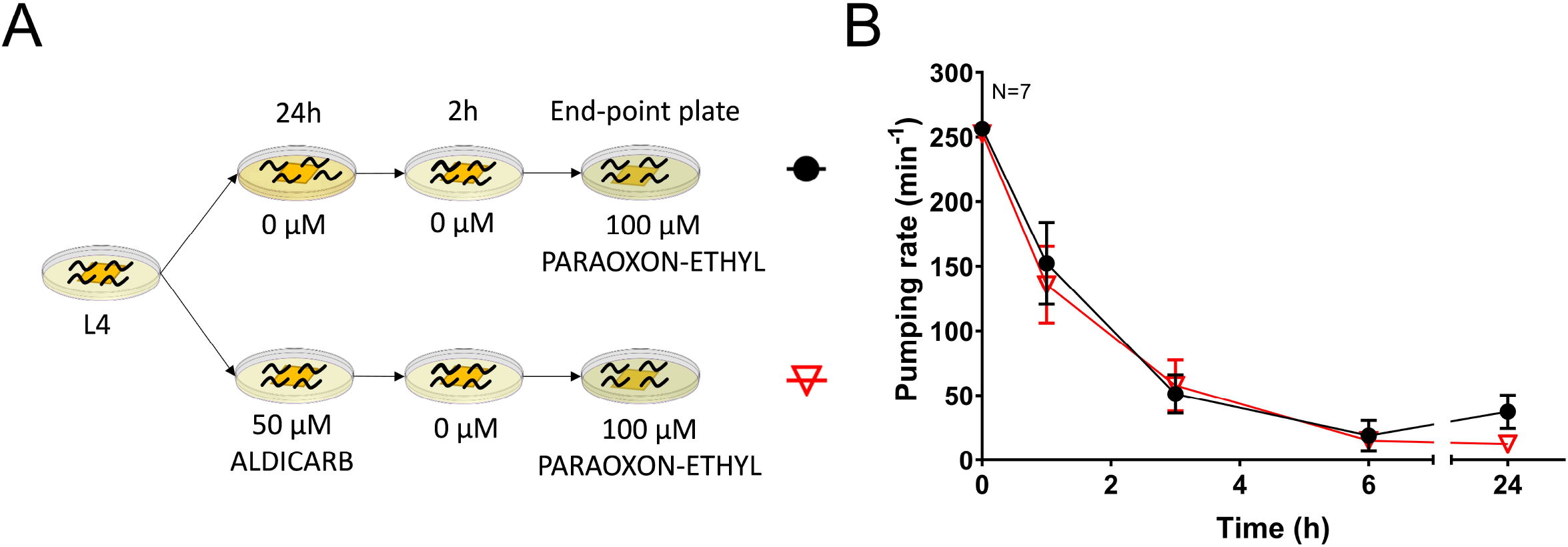
Aldicarb-preconditioned and non-preconditioned nematodes exhibit a similar pharyngeal function when exposed to maximal dose of paraoxon-ethyl. A) Synchronized L4 worms were incubated on either vehicle control or 50 μM aldicarb plates for 24 hours. After recovery on non-drug containing plates, they were exposed to a maximal dose of paraoxon-ethyl of 100 μM where pumping was measured. B) Aldicarb pre-exposed nematodes exhibit a similar sensitivity to maximal dose of paraoxon-ethyl than non-preconditioned worms. Data are shown as mean ± SEM of 7 worms in at least 4 independent experiments.

Overall, the preconditioning with aldicarb provoked the adaptation of the pharyngeal circuit to post-exposure with aldicarb but not to the post-exposure with paraoxon-ethyl. This reinforced our hypothesis that nematodes are able to adapt to anti-cholinesterase drug exposure by the previous experience. However, this is achieved by distinct mechanisms for carbamates and organophosphates. This clearly indicates that carbamates are not a well-suited model to detail investigation of the processes that underlie organophosphate intoxication, recovery and plasticity, even though they exhibit the same mode of inhibition to acetylcholinesterase (Colovic, Krstic et al. 2013).

### *C. elegans* adults exhibit spontaneous recovery of the NMJ function in the presence of higher doses of paraoxon-ethyl

A different mode of adaptation in *C. elegans* can follow upon the chronic exposure to exaggerated concentration of the stimulus (Ardiel and Rankin 2010). In the context of organophosphate poisoning, this implies a high level of acetylcholinesterase inhibition that triggers the hyperstimulation of the cholinergic receptors in the post-synaptic terminal and therefore, a pronounced effect in the cholinergic-dependent behaviours of nematodes (Izquierdo, O’Connor et al. 2020). In order to investigate how the cholinergic system of *C. elegans* responds to inhibition with exaggerated concentrations of organophosphates, we quantified the pumping rate during the sustained exposure of wild type nematodes to 500 μM paraoxon-ethyl. The concentration was calculated as 25-fold higher than the IC50 value for the pharyngeal phenotype at 24 hours and would model lethal doses that represent highly toxic environmental exposure (Izquierdo, O’Connor et al. 2020). This overstimulation of the cholinergic circuit initially caused the complete inhibition of pharyngeal pumping by 3 hours of incubation in the drug (Fig. 5A). Remarkably, after this first inhibition and despite the sustained exposure to paraoxon-ethyl, nematodes exhibited a spontaneous recovery of the pumping rate at 6 hours. This recovery of the feeding phenotype was not sustained, and the pump rate was subsequently abolished at the 24 hours of exposure to paraoxon-ethyl (Fig. 5A).

**Figure 5.**
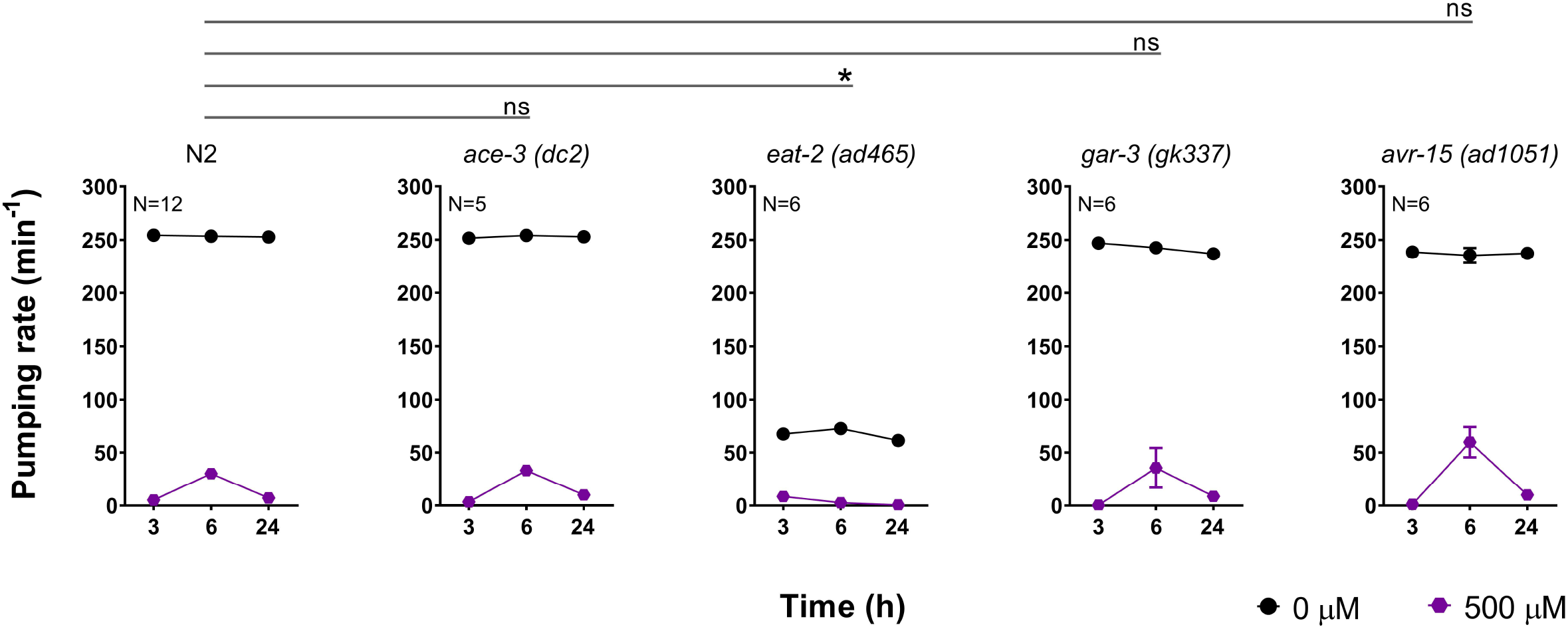
Pharyngeal and body wall neuromuscular behaviours exhibit a paraoxon-ethyl intoxication pattern characterized by three phases, an initial inhibition, a spontaneous recovery and a consequent inhibition. A) Pharyngeal pumping in nematodes exposed to 500 μM paraoxon-ethyl displays a complete inhibition of pumping at 3 hours that is spontaneously recovered at 6 hours. The complete inhibition of pumping is observed after 24 hours of exposure. Data are shown as mean ± SEM of 34 worms in at least 17 independent experiments. B) Nematodes were exposed to 500 μM paraoxon-ethyl and body length recorded 1, 3, 6 and 24 hours of exposure. Percentage of body length is referenced against the corresponding age-matched untreated control. The three phases of intoxication were observed: initial shrinkage of nematodes at 3 hours, spontaneous body length recovery after 6 hours and the subsequent shrinkage of length at 24 hours of exposure. Data are shown as mean ± SEM of 4 independent worms in 4 independent experiments. Statistical significance was calculated by two-way ANOVA test followed by Bonferroni corrections. **p≤0.01; ***p≤0.001.

The effect of the sustained exposure to 500 μM paraoxon-ethyl was additionally investigated in a different cholinergic-dependent phenotype at the level of the body wall neuromuscular junction by measuring body length. As expected, the intoxication with paraoxon-ethyl caused hypercontraction of the body wall muscles, being evidenced by the shrinkage of nematodes. Wild type worms poisoned with 500 μM paraoxon-ethyl exhibited a 50% reduction of the body length compared to non-exposed nematodes at 1 hour of incubation on drugged plates (Fig. 5B). We previously demonstrated that this reduction is the maximum level of shrinkage nematodes can reach when they are incubated to anti-cholinesterases (Izquierdo, O’Connor et al. 2020). Similar to the pharyngeal function, the body length was partially recovered at 6 hours of incubation despite the continued presence of the drug. This recovery was transitory and reversed, as the maximum shrinkage was again observed at 24 hours of exposure (Fig. 5B).

Overall, the results indicate that the overstimulation of the cholinergic pathways by high doses of paraoxon-ethyl triggers a pharyngeal and body wall behavioural response characterized by three phases: an initial inhibition, a partial and transitory spontaneous recovery and a subsequent inhibition. This highlights an adaptation process of the behaviours tested even in the continuous presence of paraoxon-ethyl. Furthermore, the identical expression of mitigating plasticity in these two phenotypes when nematodes are continuously exposed to the drug suggests a conserved mechanism underpinning the spontaneous recovery of the cholinergic function for the pumping rate and the body length.

### The molecular determinants of the pharyngeal NMJ are not involved in the cholinergic plasticity observed in the pharynx

Uncovering the signalling pathways that underpin the capacity of cholinergic-dependent behaviours to exhibit mitigating plasticity might imply the expansion of treatments that palliate aspects of organophosphate intoxication. In order to identify molecular components of this cholinergic plasticity, we compared the pumping rate of different mutant worms with the wild type control in the presence or absence of 500 μM paraoxon-ethyl at 3, 6 and 24 hours of exposure. We defined these time points as key intervals in the experiment to detect initial inhibition, spontaneous recovery and subsequent inhibition. The screening utilized the pharyngeal pumping as bio-assay since we previously demonstrated the potential of this phenotype in the research of organophosphate intoxication and recovery (Izquierdo, O’Connor et al. 2020).

We first screened the pumping rate of strains containing mutations in important components of the pharyngeal neuromuscular junction (Fig. 6). This included the acetylcholinesterase ACE-3 (Combes, Fedon et al. 2000, Combes, Fedon et al. 2003), the nicotinic receptor subunit EAT-2 (McKay, Raizen et al. 2004, Choudhary, Buxton et al. 2020), the muscarinic receptor GAR-3 (Steger and Avery 2004) and the glutamate-gated chloride channel subunit AVR-15 (Dent, Davis et al. 1997). All these proteins are critical to the contraction-relaxation cycle of the pharyngeal muscles responsible for the pumping rate. All mutants tested, except *eat-2 (ad465)*, exhibited the three phases pattern of initial inhibition of pumping, rebound recovery and reoccurring inhibition similar to the wild type control exposed to paraoxon-ethyl (Fig. 6). This indicates that these proteins are not key determinants of the drug-induced plasticity in the pharyngeal pumping phenotype.

**Figure 6.**
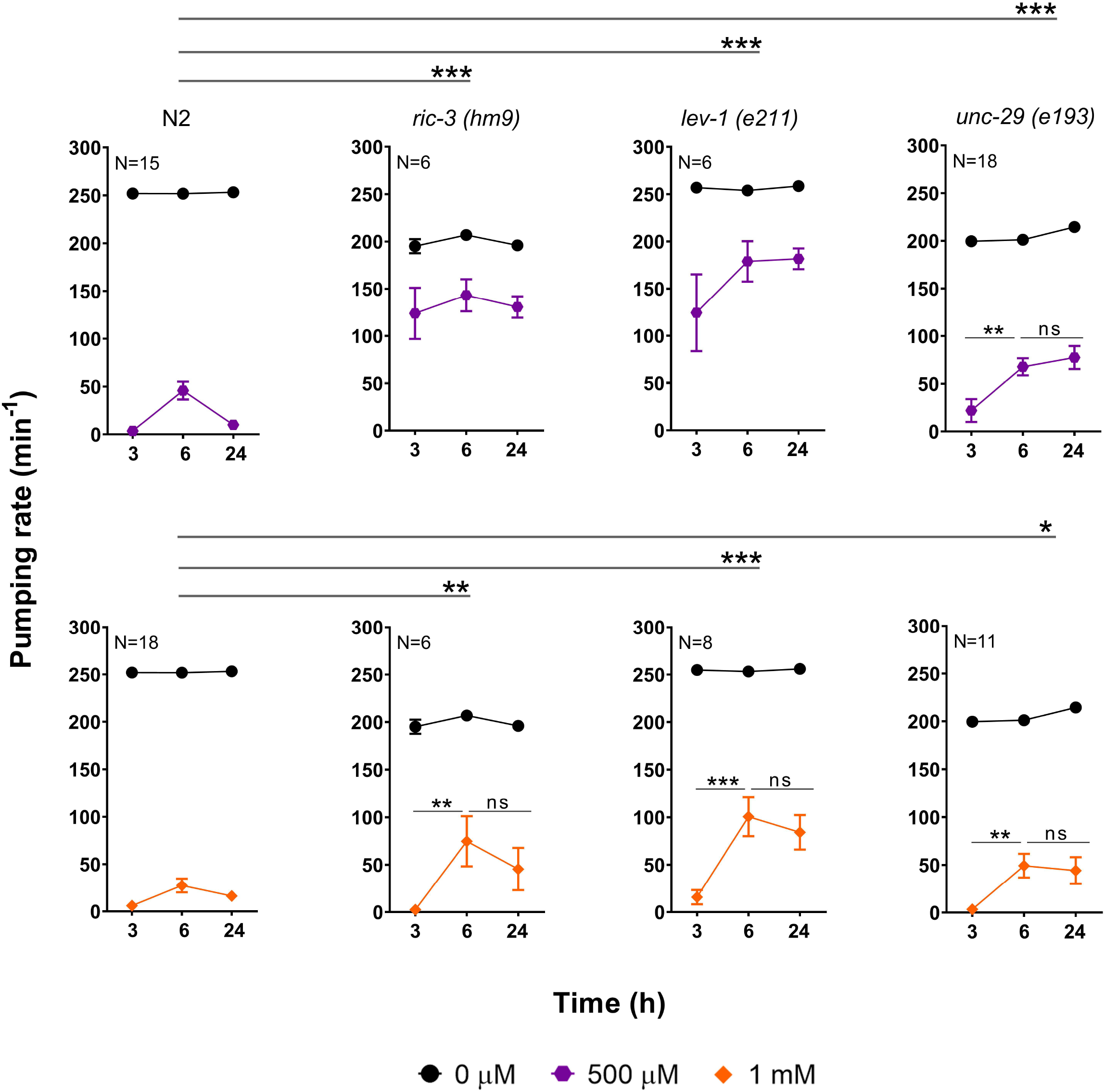
The paraoxon-ethyl induced plasticity in the pharyngeal circuit is not elicited by the NMJ components of the pharynx. N2 wild type nematodes continuously exposed to 500 μM paraoxon-ethyl exhibited spontaneous recovery of the pharyngeal function at 6 hours followed by a subsequent inhibition at 24 hours. Data are shown as mean ± SEM of 16 worms in at least 8 independent experiments. The acetylcholinesterase ACE-3 of C. *elegans* is specifically expressed in the pharyngeal muscles composing the isthmus (Combes, Fedon et al. 2003). ACE-3 deficient nematodes exposed to 500 μM paraoxon-ethyl exhibit a similar spontaneous recovery followed by inhibition of the pharyngeal pumping rate compared to wild-type worms. Data are shown as mean ± SEM of 5 worms in at least 3 independent experiments. Nematodes lacking EAT-2 did not exhibit the spontaneous recovery observed in wild type worms. Data are shown as mean ± SEM of 6 worms in at least 3 independent experiments. The muscarinic receptor GAR-3 is expressed in the isthmus and is involved in the feeding movement (Kozlova, Lotfi et al. 2019). Mutant nematodes lacking GAR-3 exhibited a similar paraoxon-induced plasticity of the pumping rate compared to the wild type worms. Data are shown as mean ± SEM of 6 worms in at least 3 independent experiments. E) *avr-15* encodes a glutamate-gated chloride channel subunit responsible for the relaxation of the pharyngeal muscle upon contraction (Dent, Davis et al. 1997). *avr-15* mutant worms exhibit a similar pattern of pharyngeal pumping rate than wild type animals intoxicated onto 500 μM paraoxon-ethyl plates. Data are shown as mean ± SEM of 6 worms in at least 3 independent experiments. Statistical significance was calculated by two-way ANOVA test followed by Bonferroni corrections. ^ns^p>0.05; *p≤0.05.

The pumping rate of *eat-2 (ad465)* deficient worms exposed to paraoxon-ethyl was continuously inhibited over the time lacking the spontaneous recovery observed in wild type nematodes at 6 hours of incubation (Fig. 6). This might hint a role in the paraoxon-induced plasticity in the feeding phenotype, but this possibility should be tempered by the intrinsic blunting of food-induced pumping. Likewise, the profound reduction of pumping rate displayed by this strain could preclude the observation of the spontaneous recovery even present.

Overall, it indicates that the spontaneous recovery observed in the pharyngeal function of nematodes exposed to paraoxon-ethyl is not determined by molecular components of the pharyngeal neuromuscular junction.

### The molecular determinants of the cholinergic plasticity in the pharynx are located in the body wall NMJ

We previously demonstrated that pharmacological activation of the body wall neuromuscular junction by either aldicarb or levamisole exerts an indirect inhibition of the pharyngeal pumping (Izquierdo, O’Connor et al. 2020). The chaperone RIC-3 and the L-type receptor subunits UNC-29 and LEV-1 are key determinants of this response and therefore are known as pharmacological determinants of the pharyngeal function (Izquierdo, O’Connor et al. 2020).

According to this, we investigated if these determinants could be involved in the paraoxon-induced plasticity observed in the feeding phenotype of wild type worms. The pumping rate of *ric-3, unc-29* and *lev-1* mutant nematodes was measured in the presence or absence of 500 μM paraoxon-ethyl at 3, 6 and 24 hours and the results were compared with the wild type control.

We observed that nematodes deficient in the ancillary protein RIC-3 and the non-alpha LEV-1 subunit of the L-type receptor exhibited a strong resistance to the pharyngeal inhibition by 500 μM paraoxon-ethyl (Fig. 7). This is consistent with our previous observations using the cholinesterase inhibitor aldicarb (Izquierdo, O’Connor et al. 2020). In addition, these mutants did not show an obvious drug-induced plasticity of the pumping rate at this concentration (Fig. 7). However, the intrinsic resistance to inhibition of the pharyngeal pumping by paraoxon-ethyl experienced by these two strains could preclude the observation or the actual expression of the drug-induced plasticity characteristic of the N2 wild type. Accordingly, we exposed nematodes deficient in *ric-3 (hm9)* or *lev-1 (e211)* to a higher dose of the cholinesterase inhibitor. The pumping rate of both strains intoxicated with 1 mM paraoxon-ethyl exhibited the initial inhibition and the spontaneous recovery phases mimicking the wild type response. However, the subsequent inhibition of the pharyngeal function observed in wild type animals is absence in *ric-3 (hm9)* and *lev-1 (e211)* strain (Fig. 7)

**Figure 7:**
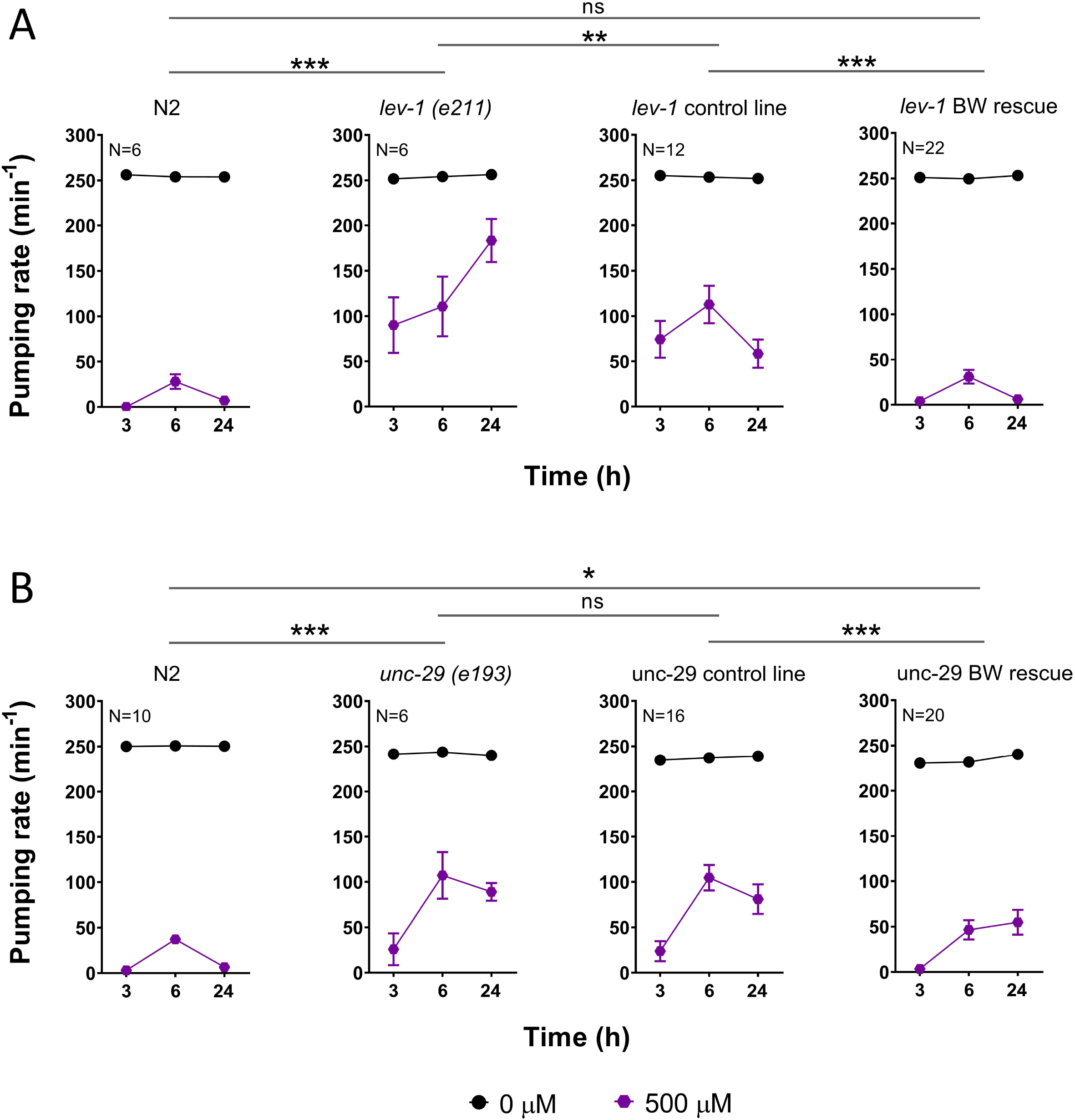
Nematodes deficient in the non-alpha subunits of the L-type body wall muscle receptor, LEV-1 and UNC-29, exhibited a sustain recovery of the pharyngeal function in paraoxon-ethyl. Paraoxon-ethyl induced plasticity of the pharyngeal function in wild type nematodes exposed to 500 μM and 1 mM. Data are shown as mean ± SEM of 15 worms in 8 independent experiments or 18 worms in 9 independent experiments, respectively. Nematodes deficient in the chaperone protein RIC-3 exhibited resistance to the inhibition of the pumping in the presence of 500 μM of paraoxon-ethyl. The exposure to 1 mM concentration inhibited the pumping rate after 3 hours and triggered a spontaneous recovery of the pharyngeal phenotype that is sustained for up to 24 hours. Data are shown as mean ± SEM of 6 worms in 3 independent experiments for each concentration. *lev-1* encodes a non-alpha subunit of the L-type receptor. LEV-1 lacking nematodes phenocopy the paraoxon-ethyl induced plasticity response of *ric-3* deficient nematodes. Data are shown as mean ± SEM of 6 worms in 3 independent experiments for 500 μM exposure or 8 worms in 4 independent experiments for 1 mM exposure. UNC-29 is the other non-alpha subunit of the L-type receptor. Nematodes deficient in UNC-29 exhibited wild type resistance to paraoxon-ethyl but sustained recovery of the pharyngeal function after 24 hours in 500 μM and 1 mM. Data are shown as mean ± SEM of 18 worms in 9 independent experiments or 11 worms in 6 independent experiments, respectively. Statistical significance was calculated by two-way ANOVA test followed by Bonferroni corrections. ^ns^p>0.05; *p≤0.05; **p≤0.01; ***p≤0.001.

Compared to wild type nematodes, both lev-1 and ric-3 mutants exhibited a sustained recovery of the pharyngeal function after the spontaneous recovery. Likewise, after the complete inhibition of the pharyngeal function after 3 hours of exposure, *ric-3 (hm9)* and *lev-1 (e211)* deficient worms displayed an exaggerated and prolonged spontaneous recovery of the pharyngeal function in the presence of the drug compared to the wild type worms (Fig. 7).

Interestingly, nematodes deficient in the other non-alpha UNC-29 subunit of the L-type receptor were not resistant to the pharyngeal inhibition by paraoxon-ethyl (Fig. 7). However, this strain presented a similar pronounced and sustained spontaneous recovery of the pharyngeal function than *lev-1 (e211)* and *ric-3 (hm9)* nematodes in the presence of 500 μM and 1 mM of paraoxon-ethyl. The three mutant strains, *ric-3 (hm9), lev-1 (e211)* and *unc-29 (e193)*, lacked the subsequent inhibition of pumping that follows the spontaneous recovery happened in the wild type worms (Fig. 7). These data show that the sustained plasticity is not a peculiarity of mutants that are insensitive to pharyngeal inhibition by paraoxon-ethyl, indicating a dissociation between the determinants of drug sensitivity and drug-induced plasticity. While RIC-3 and LEV-1 are involved in both processes, UNC-29 is only involved in the expression of the subsequent inhibition that occurs after the spontaneous recovery observed in wild type worms exposed to 500 μM paraoxon-ethyl.

In order to address this, we performed the paraoxon-ethyl intoxication experiment with transgenic lines of *lev-1 (e211)* and *unc-29 (e193)* mutant background (Fig. 8). The introduction of the wild type version of *lev-1* and *unc-29* in their respective mutant strain rescued the pharyngeal sensitivity to paraoxon-ethyl. This reinforces our previous data indicating that LEV-1 and UNC-29 are both pharmacological determinants of the drug-induced inhibition of pharyngeal function (Izquierdo, O’Connor et al. 2020). However, while the introduction of *lev-1* into CB211 strain rescued the three phases distinctive of the organophosphate-induced plasticity observed in the pharyngeal phenotype of wild type worms (Fig. 8A), the introduction of *unc-29* into CB193 strain did not (Fig. 8B). This highlights the importance of the LEV-1 subunit, and therefore the body wall L-type receptor, in the subsequent inhibition of the pharyngeal function that occurs after the spontaneous recovery in the presence of paraoxon-ethyl.

**Figure 8:**
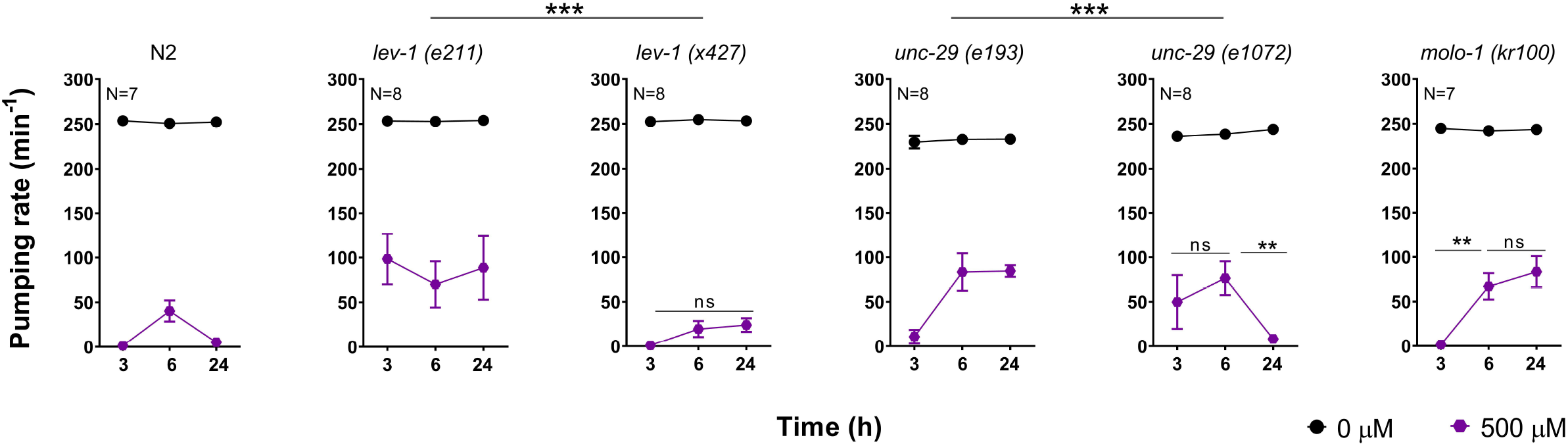
Body wall muscle rescue of the non-alpha subunits LEV-1 and UNC-29 restores wild type sensitivity to prolonged paraoxon-ethyl exposure. A) The pharyngeal sensitivity and the three phases characteristic of drug-induced plasticity to paraoxon-ethyl in the *lev-1* deficient worms were restored by introducing the wild type version of the gene selectively in the body wall muscles under control of the *myo-3* promoter. Data are shown as mean ± SEM of 6 worms in 3 independent experiments for N2 and *lev-1 (e211)* strains; 12 worms in 6 independent experiments of 2 independent lines for *lev-1* control line and 22 worms in 11 independent experiments of 4 independent lines for *lev-1* BW rescue. B) The introduction of the wild type UNC-29 in the body wall muscles of *unc-29* mutant worms rescued the consequent inhibition of the pharyngeal function that follows the spontaneous recovery in paraoxon-ethyl exposed worms. Data are shown as mean ± SEM of 10 worms in 5 independent experiments for N2; 6 worms in 3 independent experiments for *unc-29 (e193)*; 16 worms in 8 independent experiments of 3 independent lines for *unc-29* control line and 20 worms in 10 independent experiments of 3 independent lines for *unc-29* BW rescue. Statistical significance was calculated by two-way ANOVA test followed by Bonferroni corrections. ^ns^p>0.05; **p≤0.01; ***p≤0.001.

### The sensitivity of the L-type receptor at the body wall muscle is responsible of the pumping inhibition that follows the spontaneous recovery in the presence of paraoxon-ethyl

LEV-1 and UNC-29 are the two non-alpha subunits that combine with the three alpha subunits UNC-63, UNC-38 and LEV-8 to compose the L-type receptor at the body wall muscles of nematodes (Boulin, Gielen et al. 2008). However, the non-alpha subunits of these receptors create the binding pocket for neurotransmitter only when they are combined with other alpha subunit (Weise, Machold et al. 1991). To investigate the specificity of these two subunits in the paraoxon-ethyl plasticity response, we analysed the behaviour of different mutant strains in these genes in the presence of 500 μM of the cholinesterase inhibitor (Fig. 9). The *x427* allele of *lev-1* consists of 1,267 pb deletion that contains exon 4. This results in a LEV-1 protein that lacks the first, second and third transmembrane domains (Supplementary figure 1). The *e211* mutation consists of a missense substitution of glycine to glutamic acid in the fourth transmembrane domain of LEV-1 (Supplementary figure 1). Interestingly, nematodes with the *x427* allele of *lev-1* exhibited wild type sensitivity to paraoxon-ethyl and lacked the sustained recovery of the pharyngeal pumping in the presence of the drug characteristic of the strain harbouring the *e211* mutation (Fig. 7 and 9).

**Figure 9:**
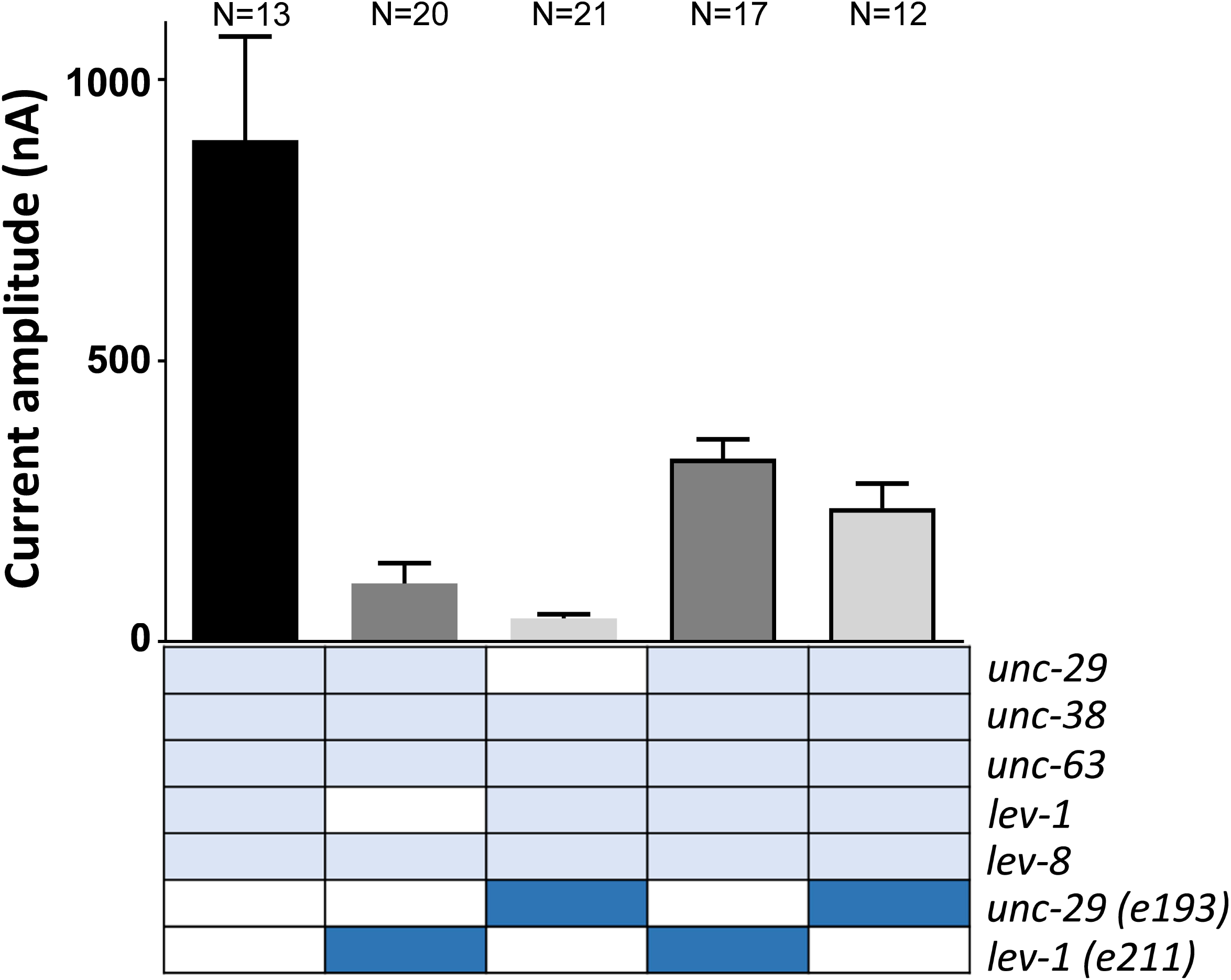
The efficacy of the L-type receptor is a significant determinant of the spontaneous recovery of pharyngeal pumping in nematodes exposed to paraoxon-ethyl. Paraoxon-induced pharyngeal plasticity in wild type nematodes incubated with 500 μM of acetylcholinesterase inhibitor. Data are shown as mean ± SEM of 14 worms in at least 7 independent experiments. *lev-1 (e211)* mutant strain contains a single point mutation in the fourth transmembrane domain. This confers resistance to paraoxon-ethyl induced inhibition of the pharyngeal pumping. Data are shown as mean ± SEM of 8 worms in at least 4 independent experiments. The mutation of *lev-1 (x427)* strain consists of a rearrangement of the genomic sequence that prevents the transcription of the gene into a protein (Fleming, Squire et al. 1997). This is the reference null-mutant of LEV-1. This mutation does not allow the expression of the mitigating plasticity in pharyngeal pumping observed in wild type worms exposed to paraoxon-ethyl. Data are mean ± SEM of 8 worms in at least 4 independent experiments. The mutation of *unc-29 (e193)* strain consists of a single point proline to serine in the loop connecting the second and third transmembrane domain of the subunit. This proline is highly conserved in the cys-loop receptor subunits and is implicated with the gating of the receptor (Connolly and Wafford 2004, Lummis, Beene et al. 2005, Albuquerque, Pereira et al. 2009). This mutation conferred a sustained paraoxon-ethyl induced plasticity. Data are shown as mean ± SEM of 8 worms in at least 4 independent experiments. In contrast, the null-mutant *unc-29 (e1072)* exhibited resistance to the pharyngeal inhibition by paraoxon-ethyl but did not express paraoxon-ethyl induced plasticity. Data are shown as mean ± SEM of 8 worms in at least 4 independent experiments. MOLO-1 is an auxiliary protein implicated in the positive modulation of the L-type receptor without affecting the location (Boulin, Rapti et al. 2012). *molo-1* lacking worms exposed to paraoxon-ethyl exhibited spontaneous recovery of the pharyngeal function that was sustained over the time compared to wild type worms. Data are shown as mean ± SEM of 7 worms in at least 4 independent experiments. Statistical significance was calculated by two-way ANOVA test followed by Bonferroni corrections. ***p≤0.001.

The strain CB1072 is considered a null mutant strain of *unc-29* (Richmond and Jorgensen 1999). The *e1072* allele contains a G to A base substitution in the splicing acceptor site of intron 8 (Supplementary figure 2). This causes a new splice acceptor that utilizes the first G in exon 9 creating a frameshift mutation and a premature stop codon between the first and the second transmembrane domain of the protein (Supplementary figure 2). However, the *e193* allele contains a missense mutation that substitutes a conserved proline for a serine in the loop connecting the second and third transmembrane domain (Supplementary figure 2). Similar to *lev-1* deficient strains, the null mutant allele of *unc-29 (e1072)* did not exhibit the sustained recovery of the pharyngeal function in the presence of paraoxon-ethyl characteristic of the *e193* allele (Fig. 9).

If we hypothesise that *lev-1 (e211)* and *unc-29 (e193)* genes harbouring non-null mutations encode for subunits that could be incorporated into the mature L-type, our data suggest that the resulting receptor might be altered in its sensitivity and/or function. This modification could be involved in the sustained recovery of the pharyngeal function after initial paraoxon-induced inhibition occurred. The fact that the paraoxon-induced plasticity in the pharyngeal phenotype observed in *lev-1 (e211)* and *unc-29 (e193)* mutant strains phenocopy the paraoxon response observed in the mutant strain *molo-1 (kr100)* supports the hypothesis (Fig. 9). Since MOLO-1 is an auxiliary protein that acts as positive modulator of the L-type receptor (Boulin, Rapti et al. 2012), the data suggest that the *lev-1 (e211)* and *unc-29 (e193)* mutations might confer different sensitivity to the L-type receptor at the body wall muscles of *C. elegans*.

To further investigate this hypothesis, the L-type of *C. elegans* was expressed in *Xenopus* oocytes by co-injecting cRNAs of the five subunits generating the ion channel (UNC-63, UNC-38, UNC-29, LEV-1 and LEV-8) along with the three ancillary proteins (UNC-50, RIC-3 and UNC-74) as previously reported (Boulin, Gielen et al. 2008). The wild type version of either LEV-1 or UNC-29 was replaced by the mutated version corresponding to the genes *e211* and *e193*, respectively, and the current amplitude to 300 μM acetylcholine was compared between the different population of receptors (Fig. 10). The co-expression of any of the mutations described along with the wild type cRNAs of the other components significantly dropped the amplitude of the current being the *e193* mutation in *unc-29* the most restrictive for the L-type receptor function (Fig. 10). However, the fact that the current amplitude is not abolished in these two subtypes of L-type receptors supports the hypothesis of the insertion of the mutated subunits in the functional ion channel. Indeed, omission to add either LEV-1 or UNC-29 cRNA in combination with the other *C. elegans* L-type receptor subunits failed to give rise to functional channels when expressed in the Xenopus oocyte (Boulin, Gielen et al. 2008). Subsequently, cRNAs of the mutated subunits, either *lev-1 (e211)* or *unc-29 (e193)*, were co-injected with the five wild type subunits in a 1:1 ratio. Interestingly, the co-expression of any of the mutations described in either *lev-1* or *unc-29* with their respective wild type cRNA significantly reduce the amplitude of the acetylcholine-elicited currents (Fig. 10). This indicates that the wild type and the mutated version of the protein compete for the formation of the mature ion channel. Such a result strongly supports a dominant-negative effect of the mutated subunit on the L-type receptor function.

**Figure 10:**
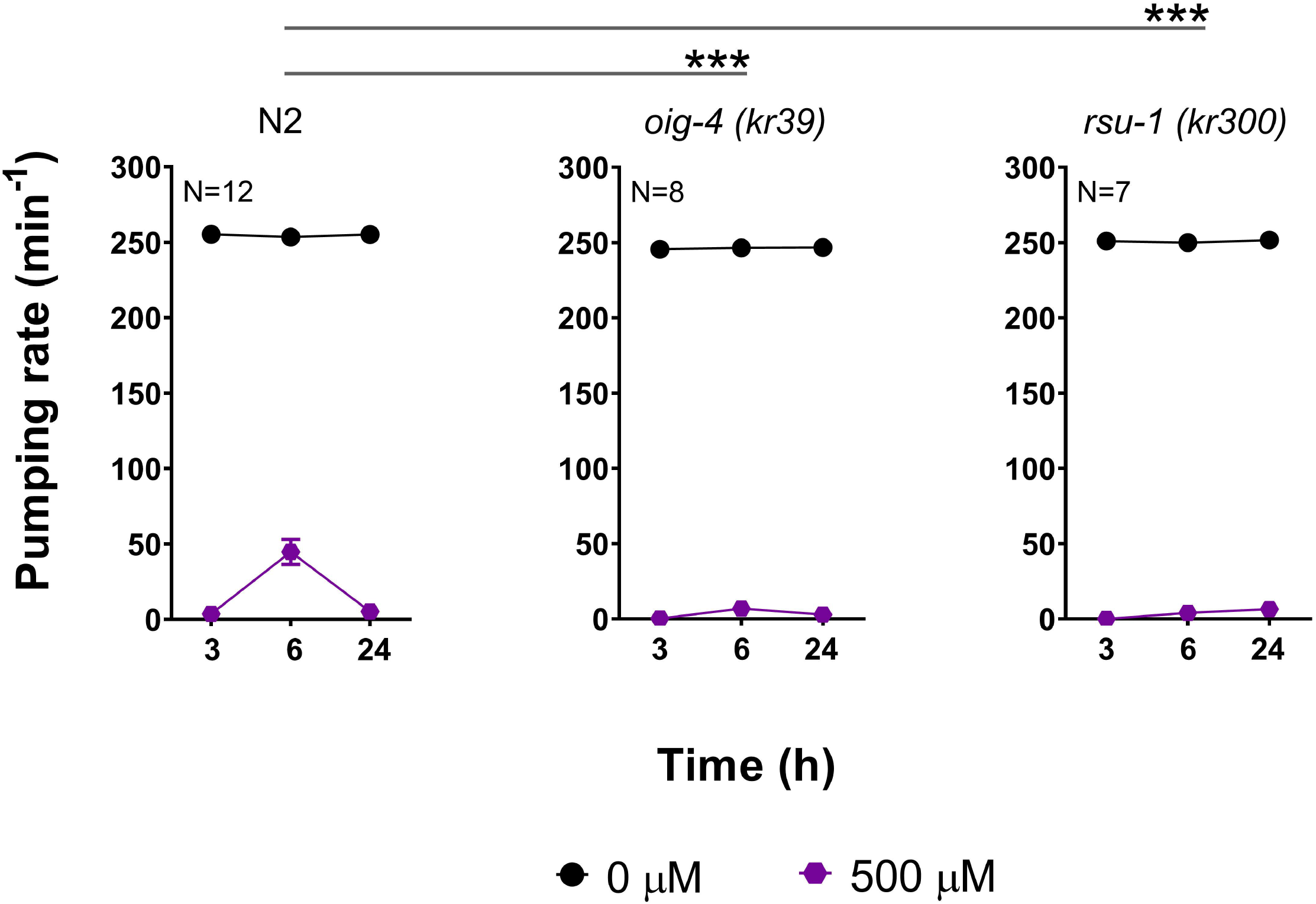
*lev-1 (e211)* and *unc-29 (e193)* encode for functional subunits that modify the signalling property of the L-type receptor. Current amplitude to 300 μM of acetylcholine was quantified in different populations of L-type receptors expressed in *Xenopus oocytes* containing either *lev-1 (e211)* or *unc-29 (e193)* mutations (dark blue). The substitution of the wild type subunits *lev-1* or *unc-29* for their respective *lev-1 (e211)* or *unc-29 (e193)* alleles reduced the current amplitude of the L-type receptor. The co-expression of wild type and mutant genes in a 1:1 ratio causes a reduction of the amplitude to acetylcholine-evoked currents, indicating both subunits compete for inclusion into the mature ion channel. Data are shown as the mean ± SEM. Numbers above bars indicate the number of oocytes recorded for each condition.

Overall, the data indicate that *lev-1 (e211)* and *unc-29 (e193)* encode for functional subunit proteins that can be inserted into the mature receptor, modifying its sensitivity to the neurotransmitter.

### The location of the L-type receptor at the body wall NMJ might be involved in the spontaneous recovery of the pharyngeal pumping in the presence of paraoxon-ethyl

Prompted by the important modulatory role of the L-type receptor in allowing the expression of mitigating plasticity to paraoxon-ethyl exposure, we tested the pharyngeal pattern to paraoxon intoxication in different auxiliary proteins of the receptor function. RSU-1 and OIG-4 are NMJ proteins that play an important role in organizing the L-type receptor within the body wall neuromuscular junction (Rapti, Richmond et al. 2011, Pierron, Pinan-Lucarre et al. 2016). These mutants exhibited wild type sensitivity to paraoxon-ethyl exposure (Fig. 11). The pharyngeal function was completely inhibited after 3 hours of incubation with the drug. However, they did not show the spontaneous recovery of the pumping rate in the presence of paraoxon as the wild type strain (Fig. 11), being both deficient in the paraoxon-induced mitigating plasticity.

**Figure 11:**
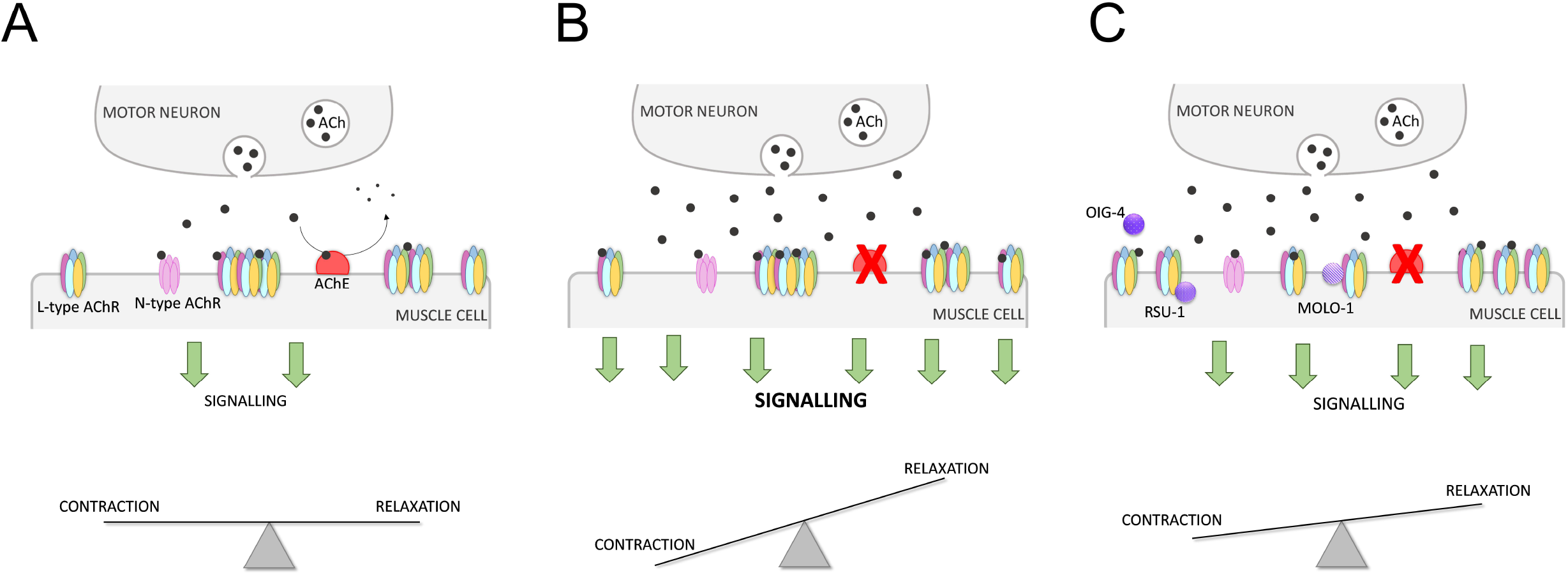
The synaptic organization of the L-type receptors underpins spontaneous recovery of the pharyngeal function observed in nematodes exposed to paraoxon-ethyl. The pharyngeal function of wild type nematodes exposed to paraoxon-ethyl exhibited drug induced plasticity. Data are shown as mean ± SEM of 12 worms in at least 6 independent experiments. OIG-4 is a muscle-secreted protein involved in the location of the L-type receptor at the body wall NMJ (Rapti, Richmond et al. 2011). *Oig-4* lacking nematodes are deficient in the spontaneous recovery of the pharyngeal pumping in the presence of paraoxon-ethyl observed in the wild type worms. Data are shown as mean ± SEM of 8 worms in at least 4 independent experiments. RSU-1 is a cytosolic muscle protein involved in maintaining the equilibrium between synaptic and extra-synaptic nicotinic receptors (Pierron, Pinan-Lucarre et al. 2016). *rsu-1* lacking nematodes exposed to paraoxon-ethyl are deficient in the spontaneously recovery of the pharyngeal function in the presence of paraoxon-ethyl. Data are shown as mean ± SEM of 7 worms in at least 4 independent experiments. Statistical significance was calculated by two-way ANOVA test followed by Bonferroni corrections. ***p≤0.001.

Overall, the data identify tuneable plasticity with respect to organophosphate intoxication which has the potential to mitigate or aggravate paraoxon-ethyl toxicity. This plasticity is driven from the function of the L-type receptor at the body wall muscle. However, distinct molecular determinants interfering in its function and/or sensitivity are additionally involved in the process.

## Discussion

Organophosphates are environmental biohazards that cause at least two million of poisoning cases and lead to an estimated 200,000 deaths annually (Jeyaratnam 1990, Eddleston and Phillips 2004, Gunnell, Eddleston et al. 2007). The nature of this intoxication is the impediment of ending the acetylcholine signal by the binding and inhibition of acetylcholinesterases in the synaptic cleft (Colovic, Krstic et al. 2013, Tattersall 2018). The overstimulation of the cholinergic transmission at the central and peripheral nervous system triggers a wide range of clinical manifestations (Jokanovic and Kosanovic 2010, Tattersall 2018). However, the pharmacological treatment to mitigate the symptoms of the cholinergic syndrome is restricted to two main mechanisms, the inhibition of muscarinic receptors by atropine and the reactivation of organophosphate-bond acetylcholinesterase by oximes. Benzodiazepines are additionally used to treat seizures during the first stage of intoxication (Eddleston and Chowdhury 2016). Since the efficiency of these medications is limited in many aspects (Buckley, Karalliedde et al. 2004, Eddleston, Buckley et al. 2004, Worek, Thiermann et al. 2004, Eddleston and Chowdhury 2016), we propose here the investigation of plasticity-promoting mechanisms in order to develop alternative pathways to mitigate against the effects of anti-cholinesterase poisoning (Fig. 12). The model organism *C. elegans* was utilized for this purpose. This free-living nematode has neuromuscular organs led by a highly conserved cholinergic pathway, compared with mammals, that triggers easy quantifiable phenotypes (Rand 2007, McVey, Mink et al. 2012, Pereira, Kratsios et al. 2015). However, these phenotypes can be modulated according present or past experiences generating homeostatic responses and plasticity (Hobert 2003, Ardiel and Rankin 2010). In previous investigations, we highlighted the quantification of pharyngeal pumping movements as the most suitable cholinergic-dependent phenotype to investigate acetylcholinesterase intoxication and recovery (Izquierdo, O’Connor et al. 2020). Here, we demonstrated that *C. elegans* nematodes are able to develop cholinergic plasticity in two different contexts, preconditioning with low doses of the drug and chronic exposure to large concentrations. However, the behavioural consequences of these two paradigms are opposite for organophosphate intoxication.

**Figure 12:**
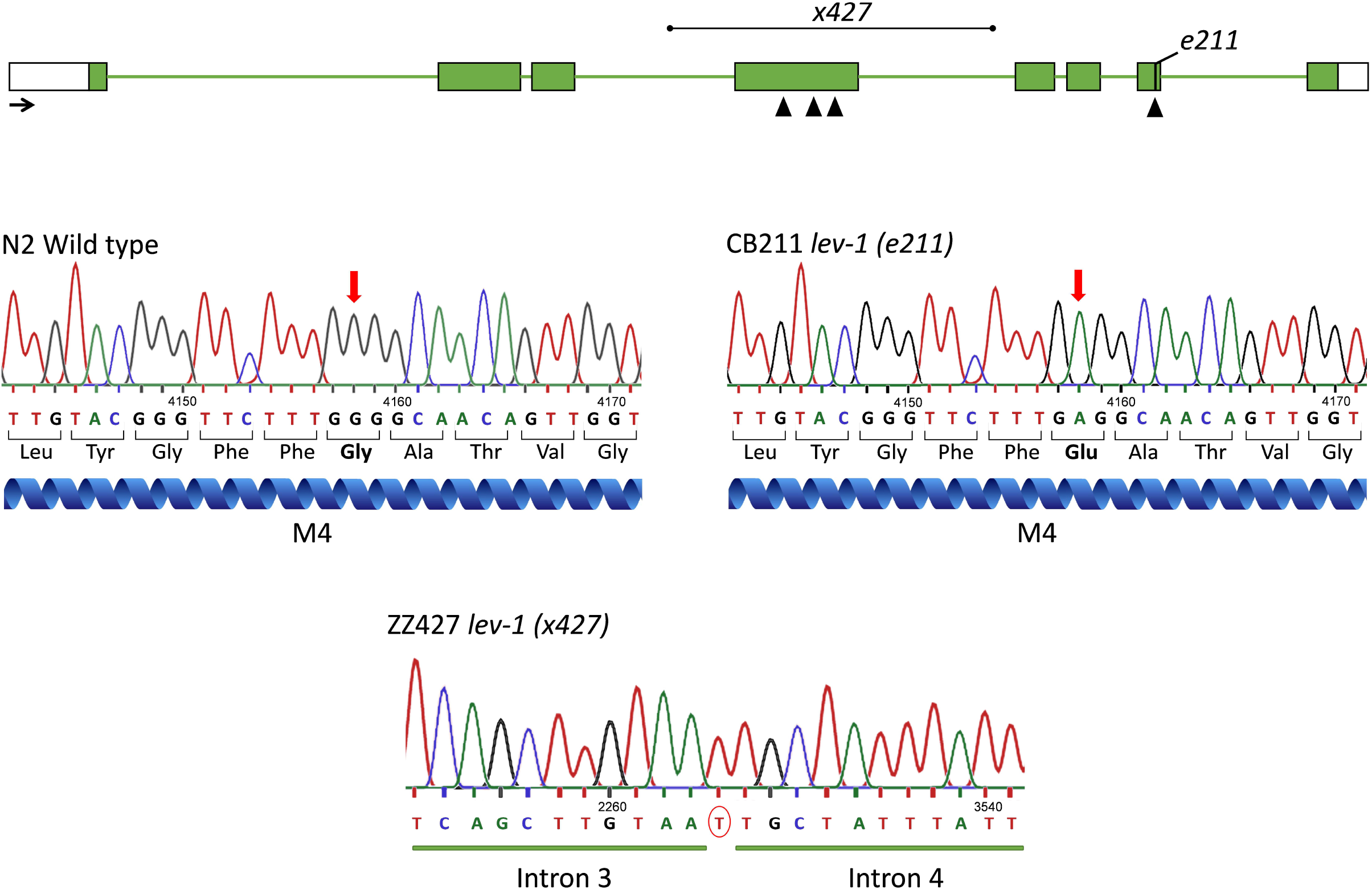
Hypothesised mechanism underpinning paraoxon-induced plasticity in nematodes exposed to 500 μM paraoxon-ethyl. A) The body wall motor neuron releases acetylcholine to the neuromuscular junction that activates L-type and N-type nicotinic receptors causing the muscle contraction. Acetylcholinesterases catalyse the breakdown of acetylcholine ending the signalling. This causes a cholinergic transmission from the motor neuron that allows balanced excitation at the body wall muscle. B) The inhibition of the acetylcholinesterase by paraoxon-ethyl causes an increase of acetylcholine at the neuromuscular junction. This leads to the overstimulation of the cholinergic receptors expressed in the muscle fibres and, therefore the hypercontraction of the muscle. C) The location of the L-type receptors at the body wall neuromuscular junction is altered in an OIG-4 and/or RSU-1 dependent mechanism. The sensitivity this L-type receptor is additionally modified in a mechanism dependent of the auxiliary protein MOLO-1. These two pathways contribute to the modulation of the cholinergic signal in the presence of paraoxon-ethyl, allowing an underlying plasticity that sustains behavioural function that is otherwise lost.

The preconditioning effect with lower doses of paraoxon-ethyl aggravated the pharyngeal inhibition in nematodes post-exposed to a larger concentration of the same drug (Fig. 3). Surprisingly, this plasticity effect with the organophosphate paraoxon-ethyl was completely opposite to the effect observed with the carbamate aldicarb in the same experiment (Fig. 2). Organophosphates and carbamates are equivalent drugs with similar mode of action (Colovic, Krstic et al. 2013). However, our data indicate that the intoxication consequences beyond the inhibition of acetylcholinesterase are different for both chemicals. This should avoid the use of one of them as model to research the other.

In the context of chronic exposure to a high concentration of the organophosphate paraoxon-ethyl, nematodes developed mitigating plasticity in the presence of the drug. This was observed by a spontaneous recovery of about 20% of the pharyngeal and body wall phenotypes after their complete initial inhibition by paraoxon-ethyl intoxication. The spontaneous recovery of the pumping rate and body length was not sustained over the time, being entirely abolished at 24 hours of exposure (Fig. 5). This indicates three phases of intoxication when nematodes are exposed to paraoxon-ethyl, the initial inhibition, the spontaneous recovery and the subsequent inhibition of pharyngeal pumping and body length.

In order to identify molecular components of this mitigating plasticity, we systematically investigated the ability to express these three phases of paraoxon-ethyl intoxication in the pharyngeal pumping of mutants involved in the cholinergic pathway. We demonstrated that cholinergic components of the pharyngeal neuromuscular junction had little effect on the expression or modulation of the plasticity observed in the wild type nematodes (Fig. 6). This builds on previous observations indicating that the *per se* inhibition of the pumping rate in the presence of the anti-cholinesterase aldicarb is mediated by determinants executing their function outside the pharyngeal system (Izquierdo, O’Connor et al. 2020). Indeed, the investigation with mutants in molecular components of the body wall neuromuscular junction highlighted the pivotal role of the L-type receptor signalling to induce the paraoxon-ethyl mitigating plasticity of the pharyngeal pumping (Fig. 12). This acetylcholine-gated cation channel is composed by the alpha subunits UNC-63, UNC-38 and LEV-8 and the non-alpha subunits UNC-29 and LEV-1 (Boulin, Gielen et al. 2008). Auxiliary and ancillary proteins have been linked with the function of the receptor by controlling trafficking, sensitivity, expression or clustering (Ramarao and Cohen 1998, Gally, Eimer et al. 2004, Boulin, Gielen et al. 2008, Gendrel, Rapti et al. 2009, Rapti, Richmond et al. 2011, Boulin, Rapti et al. 2012, Briseno-Roa and Bessereau 2014, Pierron, Pinan-Lucarre et al. 2016). We observed that *oig-4 (kr39)* and *rsu-1 (kr300)* deficient nematodes lack the spontaneous recovery of the pharyngeal pumping in the presence of paraoxon-ethyl compared with wild type worms (Fig. 11). OIG-4 is a muscle secreted protein that interacts with the complex formed by L-type receptor, LEV-9 and LEV-10 at the neuromuscular junction stabilizing the clusters (Rapti, Richmond et al. 2011). Neither *lev-9* nor *lev-10* phenocopied the paraoxon-ethyl induced plasticity observed in *oig-4* deficient animals (data not shown). However, OIG-4 remains partially localized independently of LEV-9 and LEV-10, possibly by its interaction with UNC-29, explaining the difference observed in their phenotype (Rapti, Richmond et al. 2011). RSU-1 is a protein expressed in the cytoplasm of the muscle cells and is required for the proper balance of the L-type receptor distribution between synaptic and extrasynaptic regions (Pierron, Pinan-Lucarre et al. 2016). The fact that these two proteins are involved in the location of the muscle receptor indicate that the position of the receptor during organophosphate intoxication might be altered to modulate the excess of cholinergic signal (Fig. 12). This alteration could be necessary for the expression of paraoxon-induced mitigating plasticity of the pharyngeal pumping. These data are consistent with previous observations where the distribution of nicotinic receptors at the mammalian neuromuscular junction of skeletal muscle is altered in acetylcholinesterase knockout mice to compensate the chronic absence of enzyme activity (Girard, Barbier et al. 2005).

The two non-alpha subunits, UNC-29 and LEV-1, of the L-type receptor were previously pointed as molecular determinants of the aldicarb sensitivity in the pharyngeal pumping of nematodes exposed to the carbamate (Izquierdo, O’Connor et al. 2020). Here, we observed that mutant nematodes *unc-29 (e193)* and *lev-1 (e211)* displayed a paraoxon-induced plasticity response in the pharyngeal pumping characterized by only two phases: the initial inhibition and spontaneous recovery. This spontaneous recovery is sustained over the time, lacking the subsequent inhibition observed in wild type worms (Fig. 7). Interestingly, this phenotype is not conserved in null mutant strains of *unc-29 (e1072)* or *lev-1 (x427)* but phenocopy the response observed in *molo-1 (kr100)* deficient nematodes (Fig. 9). Since MOLO-1 is an auxiliary protein involved in the sensitivity of the L-type acetylcholine receptor (Boulin, Rapti et al. 2012), the data indicate that the *e193* mutation of *unc-29* and the *e211* mutation of *lev-1* might affect the L-type receptor sensitivity or shifting at the body wall muscles influencing in its potency and/or efficiency. In addition, we demonstrated that LEV-1 and UNC-29 subunits harbouring the *e211* and *e193* mutations, respectively, are able to assemble with the alpha subunits UNC-38, UNC-63 and LEV-8, and reconstitute functional L-type receptors when expressed in *Xenopus oocytes*. The resulting L-type receptor displayed reduced response to a high concentration of acetylcholine (Fig. 10) suggesting a role of the mutated subunits as modulators of the L-type receptor activity. The fact that the *e193* mutation in UNC-29 exhibited a stronger dominant negative effect compared to *e211* in LEV-1 lays an explanation for the partial rescue of the phenotype observed in transgenic lines expressing wild type UNC-29 into the body wall muscle of a *unc-29 (e193)* mutant strain (Fig. 8B).

The *e211* mutation consists in a glycine to glutamic acid substitution at the fourth transmembrane domain of the LEV-1 subunit (Supplementary figure 1). This domain of the acetylcholine receptor subunits mediates the interaction of the receptor with the phospholipid bilayer and contributes to the kinetic of activation of the receptor. Several studies revealed that mutations in the fourth transmembrane domain of the nicotinic receptor subunits alter the channel opening and closing more than the trafficking or expression of the receptor (Li, Lee et al. 1992, Lee, Li et al. 1994, Lasalde, Tamamizu et al. 1996, OrtizMiranda, Lasalde et al. 1997, Bouzat, Roccamo et al. 1998). In fact, the equivalent mutation within the transmembrane domain of a non-alpha subunit in *Torpedo californica* reduced the potency of the receptor to high concentrations of acetylcholine but has no effect at low doses (Lasalde, Tamamizu et al. 1996). This could explain why nematodes harbouring this change exhibit wild type phenotype for locomotion and pumping but strong resistance to inhibition of these two behaviours in the presence of anti-cholinesterases (Izquierdo, O’Connor et al. 2020).

The *e193* mutation of *unc-29* consists of a proline to serine substitution at the loop connecting the second and third transmembrane domain (Supplementary figure 2). This proline is highly conserved in all the subunits of the cys-loop receptor superfamily of the ligand-gated ion channels (Deane and Lummis 2001, Mosesso, Dougherty et al. 2019). The coupling of this proline with extra cellular domains in the mammalian muscle-type receptor is essential for the gating of the channel without affecting the trafficking or expression (Lee and Sine 2005, Jha, Cadugan et al. 2007, Hanek, Lester et al. 2008, Lee, Free et al. 2009, Mosesso, Dougherty et al. 2019).

Four strains were identified by exhibiting a sustained recovery of the pharyngeal function in the presence of paraoxon-ethyl, *ric-3 (hm9), lev-1 (e211), unc-29 (e193)* and *molo-1 (kr100)*. RIC-3 is an ancillary protein responsible of the maturation of different types of acetylcholine receptors (L-type, N-type and ACR-2R) (Boulin, Gielen et al. 2008, Petrash, Philbrook et al. 2013). However, the *e211* mutation of LEV-1, the *e193* mutation of UNC-29 and MOLO-1 affect the sensitivity of the L-type receptor by decreasing the channel gating. This points that the disruption of the L-type receptor function during organophosphate intoxication might be an interesting route to mitigate the symptoms of the cholinergic syndrome (Fig. 12). The use of nicotinic receptor antagonists has been previously proposed in the treatment of organophosphate intoxication (Sheridan, Smith et al. 2005, Rosenbaum and Bird 2010, Turner, Chad et al. 2011, Amend, Niessen et al. 2020). However, this medication causes a significant hypotension due to the blockage of the cholinergic signal at the parasympathetic ganglia (Sheridan, Smith et al. 2005, Rosenbaum and Bird 2010). A more specific antagonist of the nicotinic receptor at the skeletal muscle has been also considered. In fact, there are strong evidences supporting that some oximes have a beneficial effect in the recovery from nicotinic overstimulation symptoms due to the blockage of the nicotinic receptors at the skeletal muscle (Alkondon, Rao et al. 1988, Tattersall 1993, Ring, Strom et al. 2015). However, the allosteric modulation of the nicotinic receptor sensitivity could represent the most attractive option (Sheridan, Smith et al. 2005, Rosenbaum and Bird 2010, Turner, Chad et al. 2011, Amend, Niessen et al. 2020). Negative allosteric modulators could block the nicotinic receptor activation greater as the stimulation by acetylcholine increases. Non-competitive antagonist drugs have been demonstrated to block the open of the channel *in vivo* and *in vitro* organophosphate poisoning models with beneficial effects in the recovery from intoxication (Turner, Chad et al. 2011, Seeger, Eichhorn et al. 2012, Timperley, Bird et al. 2012, Price, Docx et al. 2016).

Although further investigations will be required in order to identify pharmacological treatments to modulate the cholinergic signalling during organophosphate intoxication, our research open new insights into the mechanisms that induce cholinergic plasticity in this context (Fig. 12). To achieve this, the nematode *C. elegans* might provide an attractive *in vivo* model for screening of non-competitive nicotinic receptor antagonist with potential effects against organophosphate poisoning.

## Materials and methods

### *C. elegans* maintenance and strains

Nematodes were maintained according standard procedure (Brenner 1974). Briefly, nematodes strains were grown on NMG plates at 20°C seeded with *E. coli* OP50 as source of food. Mutant strains EN39 *oig-4 (kr39)* II, EN300 *rsu-1 (kr300)* III and EN100 *molo-1 (kr100)* III were kindly provided by Jean-Louis Bessereau Lab (Institut NeuroMyoGène, France). *ZZ427 lev-1 (x427)* IV was kindly provided by William Schafer Lab (MRC Laboratory of Molecular Biology, UK). The transgenic lines VLP1: CB211 *lev-1 (e211)* IV; *Ex[Punc-122::gfp]*; VLP2: CB211 *lev-1 (e211)* IV; *Ex[Punc-122::gfp; Pmyo-3::lev-1*] were previously available in the laboratory stock (Izquierdo, O’Connor et al. 2020). The following strains were acquired from CGC: N2 wild type, DA465 *eat-2 (ad465)* II, VC670 *gar-3 (gk337)* V, PR1300 *ace-3 (dc2)* II, MF200 *rìc-3 (hm9)* IV, CB211 *lev-1 (e211)* IV, CB193 *unc-29 (e193)* I, CB1071 *unc-29 (e1072)* I.

The following transgenic lines were generated in this work: VLP10: CB193 *unc-29 (e193)* I; Ex[*Punc-122::gfp*] VLP11: CB193 *unc-29 (e193)* I; *E×[Punc-122::gfp; Pmyo-3::unc-29*].

### Generation of *unc-29* rescue constructs

The genomic region corresponding to 3.8 kb of *unc-29 locus* was amplified using the forward and reverse primers 5′ -CAGATCTCTTATGAGGACCAACCGAC −3′ and 5′- CTCTCAAAGTCAAAAAAAGGCGAGGAG −3′ (58°C annealing temperature), respectively. The PCR product was sub-cloned into pCR8/GW/TOPO following the manufacturer protocol and subsequently cloned into pWormgate plasmid containing 2.3 kb of *myo-3* promoter (Izquierdo, O’Connor et al. 2020)

PCR amplifications were performed using Phusion High-Fidelity PCR Master Mix with HF Buffer (Thermo Fisher Scientific) following manufacturer instructions.

### Generation of transgenic lines

The marker plasmid was kindly gifted by Antonio Miranda Lab (Instituto de Biomedicina de Sevilla, Spain). It drives the expression of GFP specifically in coelomocytes of *C. elegans* (Miyabayashi, Palfreyman et al. 1999).

Control and rescue transgenic lines of *unc-29* were generated by microinjection of the corresponding plasmids into one day old adults of CB193 *unc-29 (e193)* I mutant strain (Mello, Kramer et al. 1991). A concentration of 50 ng/μl of the marker plasmid *Punc-122::gfp* was injected to generate the transgenic strain VLP10. A mixture of 50 ng/μl of *Punc-122::gfp* plasmid and 50 ng/μl of *Pmyo-3::unc-29* plasmid was microinjected to generate the transgenic strain VLP11.

The genotype of CB193 strain was authenticated by PCR amplification of the *unc-29 locus* and subsequently sequencing of the PCR product before microinjection was carried out.

### Sequencing of mutant alleles

Mutations in *lev-1* and *unc-29* mutant strains were analysed by PCR amplification (Table 1) of the corresponding genomic fragment followed by Sanger sequencing. RT-PCR was additionally performed to describe mutation in CB1072 *unc-29 (e1072)* strain following previously published protocols (Izquierdo, O’Connor et al. 2020). Briefly, a single worm from either CB1072 or N2 wild type strain was lysed and subsequently used for cDNA synthesis using SuperScript^TM^ III Reverse Transcriptase kit in a total volume of 20 μl following manufacturer protocol (Invitrogen^TM^). 5 μl of the resulting cDNA was added to a final volume of 20 μl PCR reaction with indicated oligo primers (Table 1).

**Table 1:**
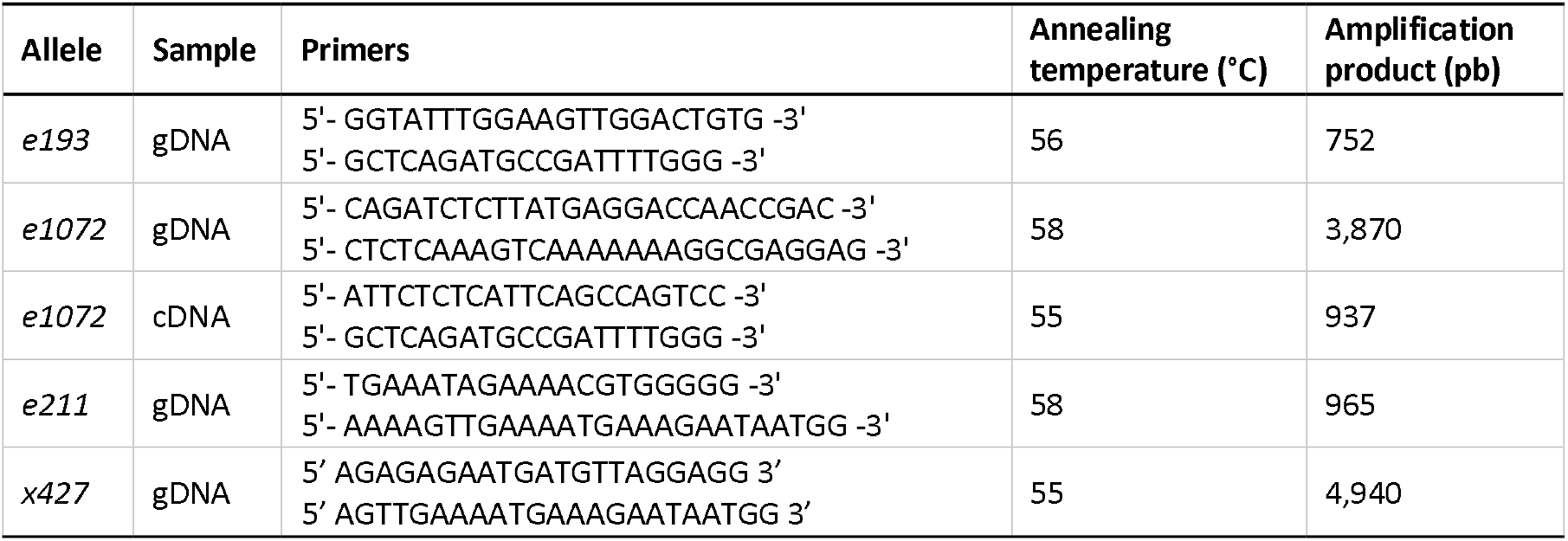
Primer sequences and PCR conditions for mutation analysis of alleles in *lev-1* and *unc-29*

PCR amplifications were performed employing Phusion High-Fidelity PCR Master Mix with HF Buffer (Thermo Scientific^™^) following manufacturer’s recommendations.

### Generation of *lev-1* and *unc-29* mutant cRNAs

*C. elegans lev-1* and *unc-29* cDNAs were cloned into the pTB207 expression vector. This vector has previously reported as suitable for transcription *in vitro* (Boulin, Gielen et al. 2008). The *e211* (G461E) and the *e193* (S258P) mutations were respectively inserted in the LEV-1 and UNC-29 subunits by PCR using the Q5 site-directed mutagenesis kit according to the manufacturer’s recommendations (New England Biolabs). The forward and reverse primers used were 5’- GTTCTTTGAGGCAACAGTTGG −3’ 5’- CCGTACAACAAAAACCGATCCA −3’ for G461E substitution in *lev-1* cDNA, and 5’ ATTCTT<u>TCA</u>CCAACATCTTCTACA −3’ / 5’- CTTTGATACAAGAAGCAAGAACAC −3’ for S258P substitution in *unc-29* cDNA. Underlined sequences indicate the mutated codons. The resulting mutant clones were sequence-checked prior linearization (Eurofins Genomics). Respective wild-type and mutant subunit cRNAs were synthesized *in vitro* with the mMessage mMachine T7 transcription kit (Invitrogen), titrated and checked for integrity. Mixes of cRNAs containing 50 ng/μL of each cRNA encoding the *C. elegans* levamisole-sensitive acetylcholine receptor (UNC-63, UNC-38, UNC-29, LEV-1 and LEV-8) subunits of interest and ancillary factors (RIC-3, UNC-50 and UNC-74) were prepared in RNase-free water (Boulin, Gielen et al. 2008).

### Oocyte electrophysiology

To investigate the functional expression of the mutated LEV-1 or UNC-29 subunits, *C. elegans* L-type nicotinic receptors, either with wild type or modified subunits, were reconstituted in *Xenopus laevis* oocytes and assayed under voltage-clamp as previously described (Blanchard, Guégnard et al. 2018). Briefly, 36 nl of cRNA mix were microinjected in defolliculated *Xenopus* oocytes (Ecocyte Bioscience) using a Nanoject II microinjector (Drummond). After 4 days incubation, BAPTA-AM-treated oocytes were voltage-clamped at a holding potential of −60 mV and electrophysiological recordings were carried out as described previously (Blanchard, Guégnard et al. 2018). Whole cell acetylcholine current responses were collected and analysed using the pCLAMP 10.4 package (Molecular Devices).

### Drug stocks

Aldicarb and paraoxon-ethyl were acquired from Merck. Aldicarb was dissolved in 70% ethanol and paraoxon-ethyl was dissolved in 100% DMSO. The drug stocks were kept at 4°C and used within one month or discarded. Obidoxime was provided by Dstl Porton Down (UK) and dissolved in distilled autoclaved water directly before use. Acetylcholine was purchased from Merk and dissolved in recording buffer (100 mM NaCl, 2.5 mM KCl, 1 mM CaCl_2_.2H_2_O, 5 mM HEPES, pH 7.3)

### Assay plates preparation

Anti-cholinesterase and obidoxime plates were prepared as previously described (Izquierdo, O’Connor et al. 2020, Izquierdo, O’Connor et al. 2020) Briefly, assay plates were made by adding a 1:1000 aliquot of the more concentrated drug stock to the molten but tempered NGM agar to obtain the indicated concentration of either aldicarb (50 μM and 250 μM) or paraoxon-ethyl (20 μM to 1 mM). 3 ml of the NGM containing the drug or the vehicle control was poured in each well of 6-well plates. After the agar solidified, plates were supplemented with 50 μl of *E. coli* OP50 (OD_600_ 1) to act as the food source. The bacterial lawn was dried on the assay plates by incubating for 1 hour in a laminar flow hood. Assay plates were finally maintained at 4°C in dark overnight. Plates were used within one day of being prepared and left at room temperature for at least 30 min before starting the experiment. There was no observable change in the bacterial lawn of drugged and control plates, therefore no effect of the anti-cholinesterase on the *E. coli* growth was discernible at any of the concentrations tested (Kudelska, Lewis et al. 2018).

The final concentration of vehicle in the drug-containing and control plates was 0.07% ethanol for aldicarb assay plates and 0.1% DMSO for paraoxon-ethyl assay plates. Neither vehicle concentrations alone affected the phenotypes tested.

### Behavioural assays

Behavioural experiments were performed at room temperature (20°C).

Pharyngeal pump rate on food was quantified by visual observation under a Nikon SMZ800 binocular microscope, using a timer and a general counter. The pumping rate was defined by the number of grinder movements per minute per worm. The pump rate was quantified for a minimum of 3 times for 1 minute each and the mean was used as pumps per minute.

The body length was measured as previously described (Mulcahy, Holden-Dye et al. 2013, Izquierdo, O’Connor et al. 2020). Briefly, images of the worms were acquired through a Hamamatsu Photonics camera and visualized for recording with IC capture software (The Imaging Source©). These images were binarized and skeletonized using ImageJ software. The length of the skeleton was used to determine the body length of the nematodes.

### Preconditioning experiments with acetylcholinesterase inhibitors

Synchronized L4 stage worms were incubated on the preconditioning plates containing either drug or vehicle control. The concentration of acetylcholinesterase inhibitor used for preconditioning was selected as one that decreased the pharyngeal pumping rate to half of the maximal response following 24 hours exposure. This was 50 μM for aldicarb and 20 μM for paraoxon-ethyl (Izquierdo, O’Connor et al. 2020). After the pre-conditioning, nematodes were transferred onto non-drug containing plates to allow the recovery of the pharyngeal function for 2 hours or 3 hours for aldicarb or paraoxon-ethyl preconditioned worms, respectively. Obidoxime plates were used during the recovery step from paraoxon-ethyl intoxication to facilitate the recovery of acetylcholinesterase after the drug inhibition (Izquierdo, O’Connor et al. 2020). Finally, nematodes were picked onto plates containing five times the concentration of drug used in the preceding preconditioning step (250 μM aldicarb or 100 μM paraoxon-ethyl). The pumping rate was measured at the indicated point times (10 min, 1, 3, 6 and 24 hours) after transferring the worms to the final control or drug treated observation plate.

### Protracted intoxication with paraoxon-ethyl

Synchronized L4 plus 1 stage nematodes were picked to either paraoxon-ethyl or vehicle control plates. Pharyngeal pump rate and body length was quantified at specified times after transferring to the assay plates (10 min, 1, 3, 6 and 24 hours). Nematodes often leave the patch of food during the first hour after transferring onto paraoxon-containing plates. They were picked back to the bacterial lawn at least 10 minutes before pump rate was measured.

### Statistical analysis

The collection of data was performed blind, viz, the experimenter was unaware of the genotype tested in each trial.

Data were analysed using GraphPad Prism 8 and are displayed as mean ± SEM. Statistical significance was assessed using two-way ANOVA followed by post hoc analysis with Bonferroni corrections where applicable. This post hoc test was selected among others to avoid false positives. The sample size N of each experiment is specified in the corresponding figure.

## Acknowledgements

We thank Dr Jean-Louis Bessereau, Laure Granger, Dr Denise Walker and Dr William Schafer for sharing strains; Dr Antonio Miranda-Vizuete for sharing *Punc-122::gfp* marker plasmid.

Additional *C. elegans* strains were provided by the CGC, which is funded by NIH Office of Research Infrastructure Programs (P40 OD010440).

## Author contributions

**Patricia G. Izquierdo:** Conceptualization, Data curation, Formal analysis, Investigation, Methodology, Validation, Visualization, Roles/Writing - original draft, review & editing. **Claude L. Charvet:** Data curation, Formal analysis, Investigation, Methodology, Writing – review & editing. **Cedric Neveu:** Funding acquisition, Writing - review & editing. **Vincent O’Connor:** Conceptualization, Funding acquisition, Methodology, Supervision, Writing - review & editing. **Christopher Green:** Conceptualization, Funding acquisition, Methodology, Supervision, Writing - review & editing. **Lindy Holden-Dye:** Conceptualization, Funding acquisition, Methodology, Supervision, Writing - review & editing. **John Tattersall:** Conceptualization, Funding acquisition, Methodology, Supervision, Writing - review & editing.

---

**Supplementary figure 1:**
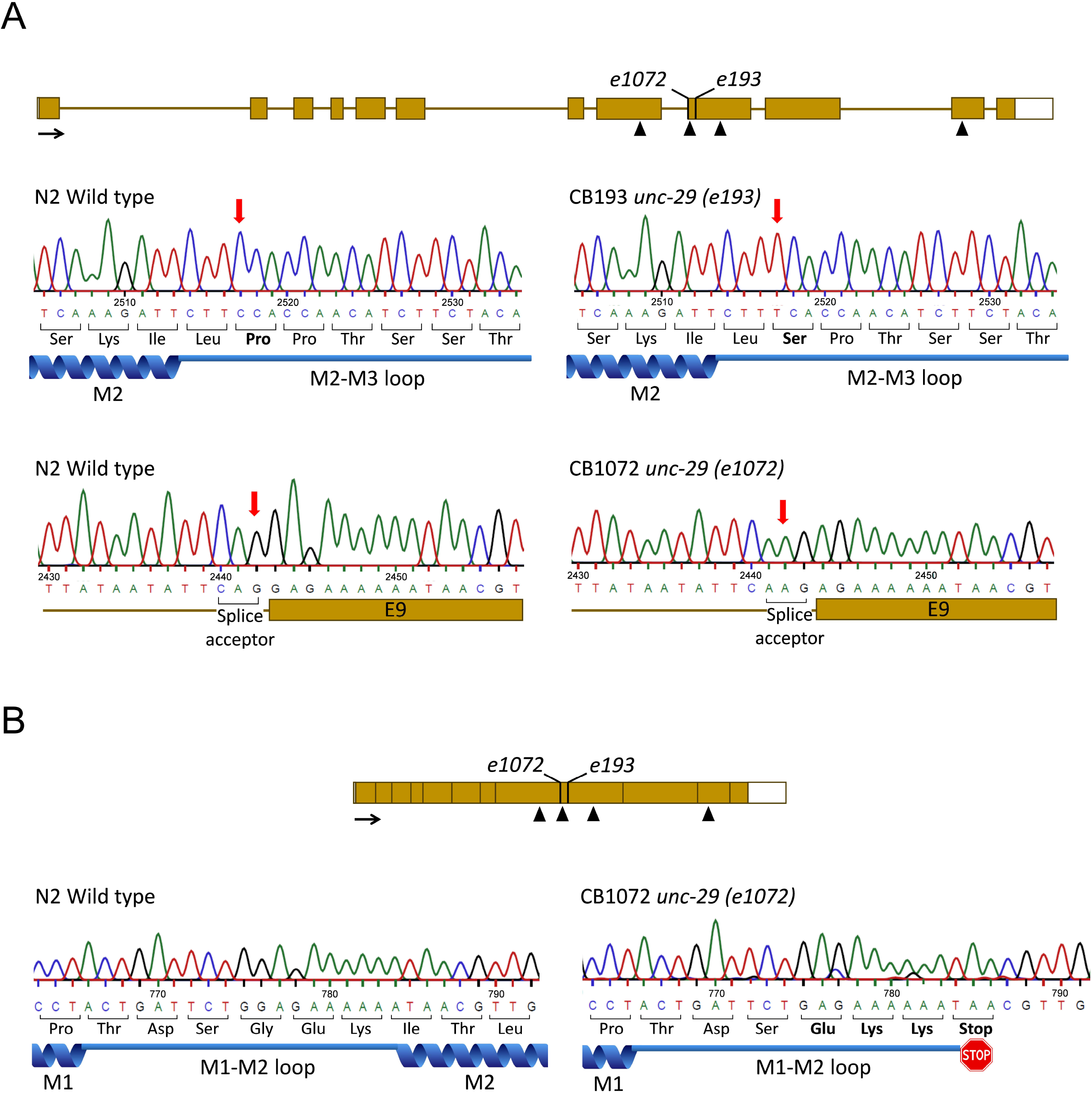
*lev-1* mutant alleles of strains CB211 and ZZ427. Genomic organization of *lev-1 locus* indicating the position of the single point mutation in *e211* and the deletion in *x427* alleles. CB211 *lev-1 (e211)* mutant strain contained a G to A missense mutation identified in exon 7 of the genomic DNA. This leads to a glycine to glutamic acid exchange at the fourth transmembrane domain (M4). The strain ZZ427 *lev-1 (x427)* contains a deletion of 1,267 bp from intron 3 to intron 4 and an T insertion. This causes a LEV-1 protein lacking the first, second and third transmembrane domain. Black arrow represents 100 pb and the sense of transcription. Black triangles in the genomic DNA represents the position of the four transmembrane domains. Chromatograms corresponds to the 5’ to 3’ readout of the minus strand. The position indicated in each chromatogram corresponds to the position of the respective base from the ATG starting codon in the genomic DNA of N2 wild type.

**Supplementary figure 2:**
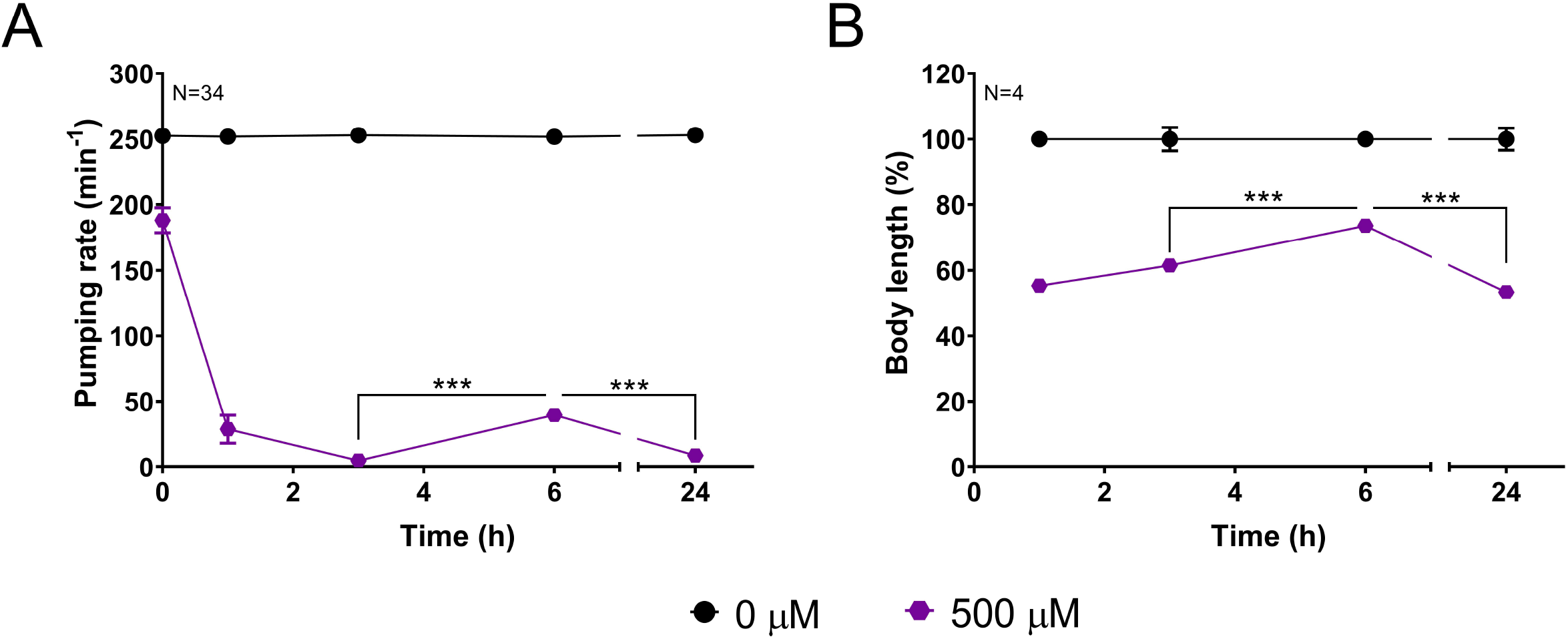
*unc-29* mutant alleles of strains CB193 and CB1072. A) Genomic organization of *unc-29 locus* indicating the position of the single point mutation in *e193* and *e1072* alleles. CB193 *unc-29 (e193)* mutant strain contained a C to T missense mutation identified in exon 9 of the genomic DNA. This causes the exchange of conserved proline to serine at a conserved residue within the extracellular loop that connects the second (M2) and third (M3) transmembrane domain of the unc-29 subunit. The strain CB1072 *unc-29 (e1072)* contains a G to A single point mutation identified in the splicing acceptor of intron 8 in the genomic DNA. This caused the formation of a new splicing site utilizing the first G in exon 9 (E9). B) RNA organization of *unc-29* indicating the position of the single point mutation in *e193* and *e1072* alleles. The mutation of *e1072* allele causes frameshift leading to a premature stop codon at the end of the predicted intracellular loop that connects the first (M1) and second (M2) transmembrane domains. Black arrow represents 100 pb and the sense of transcription. Black triangles in the genomic and cDNA represents the position of the four transmembrane domains. The position indicated in each chromatogram corresponds to the position of the respective base from the ATG starting codon in the genomic (A) or cDNA (B) of N2 wild type.

